# Heterogeneity in Functional Connectivity: Dimensional Predictors of Individual Variability during Rest and Task fMRI in Psychosis

**DOI:** 10.1101/2023.10.04.560971

**Authors:** Maria T. Secara, Lindsay D. Oliver, Julia Gallucci, Erin W. Dickie, George Foussias, James Gold, Anil K. Malhotra, Robert W. Buchanan, Aristotle N. Voineskos, Colin Hawco

## Abstract

**Background:** Individuals with schizophrenia spectrum disorders (SSD) often demonstrate cognitive impairments, associated with poor functional outcomes. While neurobiological heterogeneity has posed challenges when examining social cognition in SSD, it provides a unique opportunity to explore brain-behavior relationships. We examined the relationship between behavioral data and individual variability of functional connectivity at rest and during an emotional-processing task.

**Methods:** Neuroimaging and behavioral data were analyzed for 193 individuals with SSD and 155 controls (total n = 348). Individual variability was quantified through mean correlational distance (MCD) of functional connectivity between participants; MCD was defined as a global ‘variability score’. Hierarchical regressions were performed on variability scores derived from resting state and Empathic Accuracy (EA) task functional connectivity data to determine potential predictors (e.g., age, sex, neurocognitive and social cognitive scores) of individual variability.

**Results:** SSD showed greater MCD during rest (p = 0.00013) and task (p = 0.022). In the hierarchical regression, diagnosis remained significant when social cognition was included during rest (p = 0.008), but not during task (p = 0.50); social cognition was significant during both rest and task (both p = 0.01).

**Conclusions:** Diagnostic differences were more prevalent during unconstrained resting scans, whereas the task pushed participants into a more common pattern which better emphasized transdiagnostic differences in cognitive abilities. Focusing on variability may provide new opportunities for interventions targeting specific cognitive impairments to improve functional outcomes.

## 1. Introduction

Schizophrenia spectrum disorders (SSD) are characterized by remarkable interindividual variability in clinical symptoms (Tsuang, Lyons, and Faraone 1990), treatment response (Malhotra 2015), prognosis (Huber 1997), and pronounced cognitive and neurobiological heterogeneity (Van Rheenen et al. 2017; Viviano et al. 2018; Hawco et al. 2019). Cognitive deficits represent a core feature of SSD (Mesholam-Gately et al. 2009; Green, Horan, and Lee 2015); they are associated with functional outcomes (Nuechterlein et al. 2008), and with social cognitive impairments mediating the relationship between real-world functioning and neurocognition (Fett et al. 2011; Oliver et al. 2019; Schmidt, Mueller, and Roder 2011). Most studies thus far have treated variability as noise, and our understanding of the neurobiology of SSD has been constrained by case-control study designs that fail to consider within-group heterogeneity (Carruthers et al. 2019). However, individual variability is an important by-product of meaningful individual differences (MacDonald, Nyberg, and Bäckman 2006; Van Horn, Grafton, and Miller 2008).

Recently, our group has examined individual variability of brain activation (Hawco et al. 2020; Gallucci, Tan, et al. 2022; Gallucci, Pomarol-Clotet, et al. 2022). Individuals with SSD have exhibited greater overall individual variability than typically developing controls (TDC) in functional brain activity during cognitive tasks, with more idiosyncratic patterns related to illness duration (Gallucci, Pomarol-Clotet, et al. 2022; Gallucci, Tan, et al. 2022). The range and heterogeneity of behavior and neural circuit activation in SSD frequently exhibits overlap with behavior and activation patterns observed in TDC and may account for the inconsistencies in case-control findings (Oliver et al. 2021; Hawco et al. 2019). Previous findings have shown individuals with SSD make use of the same neural networks as TDC but generally, though not always, fall into the ‘poor performing’ side of the spectrum (Sheffield et al. 2017; Oliver et al. 2021). Examining cases and controls with a dimensional as opposed to categorical approach may lead to important insights into common and distinct neural circuits and better consider heterogeneity in clinical samples (Hajdúk et al. 2018; Hawco et al. 2019; Gallucci, Tan, et al. 2022).

Individual brains are characterized by underlying intrinsic functional connectivity profiles that are unique to the individual (Finn et al. 2015). Functional brain systems are defined by this underlying network architecture (Yeo et al. 2011; Ji et al. 2019), which is modified as necessary in response to cognitive demands (Cole et al. 2014; Mennes et al. 2010). While the majority of the literature on functional connectivity has focused on ‘task-free’ resting state fMRI (Smith et al. 2013; Yang, Gohel, and Vachha 2020), examining connectivity during task states provides additional information on the relationship between connectivity and cognition (Cole et al. 2021; Finn and Bandettini 2021; Stevens 2016; Gratton et al. 2018; Ito et al. 2020). A prior study from our group has shown that participants with SSDs fall along a similar spectrum as controls in network connectivity during a social cognitive task (Oliver et al. 2021), while another found that resting state connectivity may be more predictive of social and non-social cognitive outcomes than task-based connectivity (Viviano et al. 2018). This suggests task and resting state connectivity may provide different information regarding the relationship between cognition, functional networks, and diagnostic groups (Finn 2021).

The aim of this study was to investigate the relationship between individual variability in functional connectivity during resting state and the performance of a social task and social and non-social cognition in a large sample of TDC and individuals diagnosed with SSD. We hypothesised that individuals with SSD would display greater heterogeneity in functional connectivity than TDC, and that worse cognitive performance would be associated with greater individual variability (Gallucci, Tan, et al. 2022). The emotional-processing fMRI task will capture variability that pertains to higher-level social cognitive circuitry, while the resting state will represent more unconstrained functional connectivity variability. We used hierarchical regression analysis to examine predictors of individual variability (e.g., age, sex, diagnostic group, and neurocognitive and social cognitive scores) dimensionally across individuals with SSD and TDC.

## 2. Methods and Materials

### 2.1 Participants

Previously collected data was analyzed from the National Institute of Mental Health (NIMH)-funded “Social Processes Initiative in the Neurobiology of Schizophrenia(s) (SPINS)” multicenter study, which, in the context of the Research Domain Criteria (RDoC) framework, investigates the clinical, behavioral, and neural correlates of social cognition in SSD (Viviano et al. 2018; Oliver et al. 2021; Hawco et al. 2019). Recruitment occurred at three sites: the Centre for Addiction and Mental Health (Toronto), Zucker Hillside Hospital (New York), and the Maryland Psychiatric Research Center (Maryland) from 2014 to 2020. Participants with SSD met DSM-5 criteria for schizophrenia, schizoaffective disorder, schizophreniform disorder, delusional disorder, or psychotic disorder not otherwise specified, assessed using the Structured Clinical Interview for DSM-IV-TR (First et al. 2002). Information regarding participant medication was collected, and chlorpromazine (CPZ) equivalents were calculated for the SSD group (Leucht et al. 2016). Individuals with SSD were symptomatically stable, and had no change in antipsychotic medication or decrement in functioning/support level during the 30 days prior to enrollment. TDC did not have a current or past Axis I psychiatric disorder, excepting adjustment disorder, phobic disorder, and past major depressive disorder (over two years prior; presently unmedicated), or a first degree relative with a history of psychotic mental disorder.

Additional exclusion criteria included a history of head trauma resulting in unconsciousness, a substance use disorder (confirmed by urine toxicology screening), intellectual disability, debilitating or unstable medical illness, or other neurological diseases. All participants provided written informed consent and the protocol was approved by the respective research ethics and institutional review boards. All research was conducted in accordance with the Declaration of Helsinki.

### 2.2 Clinical and Behavioral Assessments

Psychiatric symptoms were evaluated in the SSD sample using the Brief Psychiatric Rating Scale (BPRS; (Overall and Gorham 1962)) and an adapted version of the Scale for the Assessment of Negative Symptoms (SANS; (Buchanan et al. 2007; Andreasen 1982)). The Birchwood Social Functioning Scale (BSFS; (Birchwood et al. 1990)) was administered to evaluate social functioning (omitting Employment) and the Wechsler Test of Adult Reading (WTAR; (Wechsler 2001)) as a measure of premorbid IQ.

Cognitive assessments consisted of social cognitive measures and a neurocognitive battery. Out-of-scanner social cognitive measures included the Penn Emotion Recognition Test (ER40; (Kohler et al. 2000)), the Reading the Mind in the Eyes Test (RMET; (Baron-Cohen et al. 2001)), and the Awareness of Social Inference Test–Revised (TASIT), parts 1, 2, and 3 (McDonald, Flanagan, and Rollins 2011). A ‘mentalizing’ factor score representing higher level social cognition (e.g., sarcasm detection, theory of mind) was computed as previously described (Oliver et al. 2019), from subscales of the TASIT; specifically the TASIT 2 simple sarcasm, TASIT 2 paradoxical sarcasm, and TASIT 3 sarcasm scores. While this factor analysis also identified a lower level ‘simulation’ factor, the two factors were highly correlated, and mentalizing is preferentially associated with functional outcomes and negative symptoms (Oliver et al. 2019). Thus, only the mentalizing factor score was utilized in this study. The self-report empathic concern subscale of the Interpersonal Reactivity Index (IRI; (Davis 1983)) was included as a metric for emotional empathy, which is distinct from cognitive empathy captured by the mentalizing score (Davis 1983; Shamay-Tsoory, Aharon-Peretz, and Perry 2009). Finally, neurocognition was assessed using the Measurement and Treatment Research to Improve Cognition in Schizophrenia (MATRICS) Consensus Cognitive Battery (MCCB, (Nuechterlein et al. 2008)).

### 2.3 Empathic Accuracy (EA) Task

The Empathic Accuracy (EA) task (Zaki et al. 2009) was completed during fMRI (Kern et al. 2013; Olbert et al. 2013). Participants watched 9 short videos (average length = 2.05 min, range 2.00 to 2.50 mins), presented in three runs. During each video, actors describe autobiographical events with a range of emotional valence while the participants provide continuous ratings of how positive or negative the individuals in the video are feeling (ranging from 1 (very negative) to 9 (very positive) using a button box). An EA score was calculated for each participant by correlating their ratings with self-ratings provided by the actors in the videos, providing a measure of interpersonal understanding (Zaki et al. 2009). A control condition was presented between videos, twice per run (40 s each), in which participants provide continuous ratings of the relative light or darkness of a grayscale circle as it changes shades. This was included to ensure that participants are engaged in the task and comprehend it. See the Supplement for additional details.

### 2.4 MRI Data Acquisition

Scanning was conducted using harmonised scanning parameters on six 3T scanners (see Supplementary Material). The EA task was part of a longer multimodal MRI protocol, as previously described (Viviano et al. 2018). Three EA fMRI runs were acquired using an echo-planar imaging (EPI) sequence (TR = 2000 ms, TE = 30 ms, flip angle = 77°, field of view = 218 mm, in-plane resolution = 3.4 mm2, slice thickness = 4 mm). One seven-minute resting state scan was acquired using an accelerated EPI sequence (TR = 2000 ms, TE = 30 ms, flip angle = 77°, field of view = 200 mm, in-plane resolution = 3.125 mm2, slice thickness = 4 mm, no slice gap) where participants were instructed to let their mind wander with their eyes closed for the duration of the scan. Anatomical T1-weighted scans were collected using a fast-gradient sequence (CMH and ZHH used a BRAVO sequence in sagittal plane with TR = 6.4/6.7 ms, TE = 2.8/3 ms, flip angle = 8°, voxel-size =0.9 mm; CMP, MRC, MRP and ZHP used a MPRAGE sequence in sagittal plane with TR =2300 ms, TE = 2.9 ms, flip angle = 9°, voxel-size = 0.9mm) for use in the preprocessing pipeline.

All scans underwent quality control using an in-house quality control system dashboard (https://github.com/TIGRLab/dashboard). Participants were excluded for excessive motion (average framewise displacement (FD) > 0.5 mm) and EA or control task performance indicative of disengagement or a lack of comprehension (see Figure 2 and Supplementary Material).

### 2.5 fMRI Preprocessing

Scans were preprocessed using fMRIPrep 1.5.8 (Esteban et al. 2019), based on Nipype 1.4.1 (K. J. Gorgolewski et al. 2018). Full details of the preprocessing pipeline are presented in the supplemental materials. In brief, fMRI data underwent fieldmapless distortion correction, were realigned for motion, and slice time corrected. Data was then registered to the cortical surface in the Human Connectome CIFTI space by coregistering to the T1 scan following tissue segmentation in freesurfer and nonlinear registration (Dickie et al. 2019). Scans were surface smoothed at 2mm. For both resting state and EA data, the nuisance regression model included regressors for the six head motion correction parameters, mean white matter signal, mean cerebral spinal fluid signal, the square, derivative, and square of the derivative for each of these regressors, and global brain signal (generated by fMRIPrep) (Muschelli et al. 2014; Satterthwaite et al. 2013). For the EA task, an amplitude-modulated general linear model was performed using AFNI’s 3dDeconvolve module in Nipype (Abraham et al. 2014; Cox and Hyde 1997; K. Gorgolewski et al. 2011). To model the stimulus-evoked response, the regressors were fit to each voxel and the residual activation was retained for background connectivity analysis (see Supplementary Material; (Oliver et al. 2021; Al-Aidroos, Said, and Turk-Browne 2012)). Background connectivity allows an examination of state-related connectivity rather than stimulus-driven coactivation (Al-Aidroos, Said, and Turk-Browne 2012; Norman-Haignere et al. 2012).

### 2.6 Connectivity Matrix Construction

The average time series was extracted from MNI-space fMRI data using the 360 cortical region surface based Multimodal Parcellation (MMP) 1.0 atlas (Glasser et al. 2016; Ji et al. 2019). The Melbourne Subcortex Atlas (Scale II) (Tian et al. 2020) was used to extract 32 subcortical regions. A mean time series was extracted for each of the 392 regions for both resting state and EA residual data. Only time points during the EA videos were retained. Functional connectivity matrices for each participant were generated by calculating a Fisher z transform of the Pearson correlation coefficients between each pair of regions to generate a 392 x 392 connectivity matrix, separately for rest and EA.

### 2.7 Dimensionality Reduction

Data was analyzed using RStudio v1.4.1717. The dimensionality reduction was used for both resting state and EA task fMRI data. To address the motion confound in the EA task, edge-wise correlations with FD were calculated and connectivity values that had a correlation of greater than ± 0.30 with FD were removed from the analysis (10,454 edges removed). This allowed the motion effects to be thoroughly removed from the EA task data while preserving relationships with other behavioral variables (see Supplementary table S1). This step was not performed for the resting state data as no connectivity values had a correlation of greater than ± 0.30 with FD. The upper triangle of each matrix was extracted and vectorized to provide 76,636 unique connectivity values per participant for resting state and 66,182 connectivity values per participant for the EA task. Since we were interested in investigating individual variability, our analysis of connectivity data was limited to the top ten percent of most variable connections (Xia et al. 2018). This was measured by median absolute deviation (MAD), which has been proven to be more robust against outliers than standard deviation (Xia et al. 2018). MAD was used for dimensionality reduction; the final input data consisted of 7664 unique connections for rest and 6618 connections for the EA task (Figure 1).

**Figure 1.**
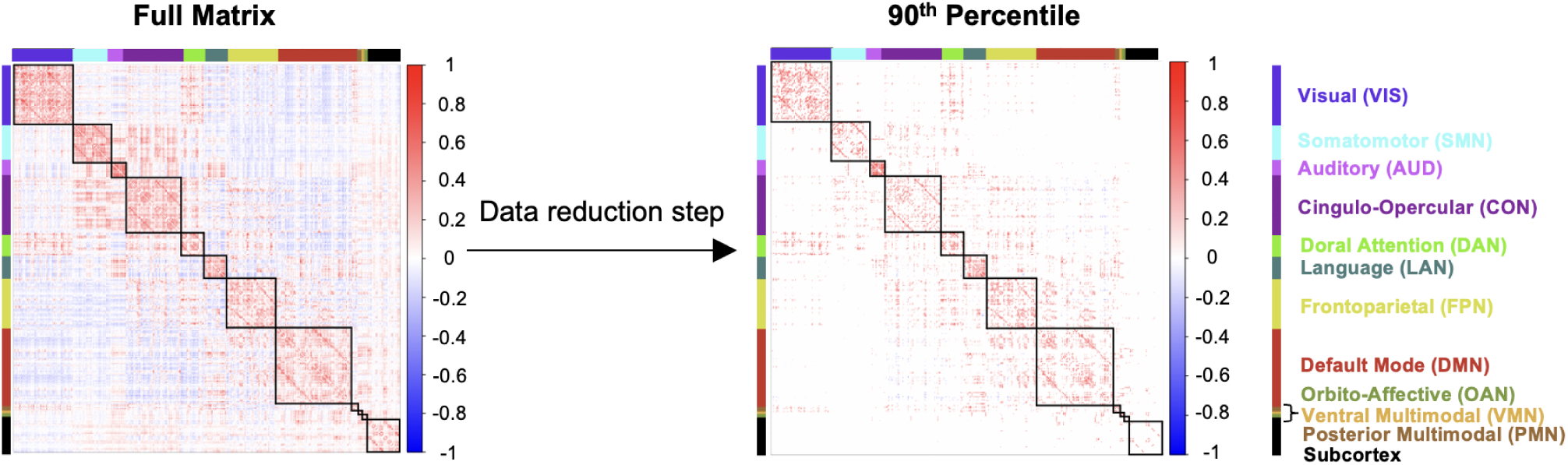
Matrix Dimensionality Reduction. Representative examples of connectivity feature selection using median absolute deviation, defined as the median of the difference between each element and the median in a vector. From the full connectivity matrix (left image), the top 10% (7664 for resting state; 6618 for EA task) of connectivity features that were the most variable across the sample were selected as a data reduction method (right matrix), and used for further analysis.

### 2.8 Individual Variability in Connectivity via Correlational Distance

The described approach was used for both resting state and EA task fMRI data. Each participant’s connectivity values were combined into a matrix (Resting state = participants x 7664 connectivity values; EA task = participants x 6618 connectivity values). Harmonisation of multi-site imaging data was accomplished using the neuroCombat package in RStudio, to account for scanner effects using an empirical Bayes framework (Johnson, Li, and Rabinovic 2007; Fortin et al. 2018; Yu et al. 2018). Age, sex, diagnostic group, framewise displacement, and clinical and behavioral scores were included in a design matrix specifying covariates, to ensure that variance related to those measures was preserved during data harmonization. A correlational distance matrix was generated, measuring the dissimilarity in connectivity across each pair of participants. The mean pairwise correlation distance for each participant to all other participants represents individual variability - a single value used as a metric to evaluate how similar or different each participant is compared to the entire group. Lower mean correlational distances indicated participants with connectivity patterns that were more similar to the overall group, and higher mean correlational distances indicated more idiosyncratic connectivity (Hawco et al. 2020; Gallucci, Tan, et al. 2022). Each participant in the study received mean correlational distance values for their EA task background connectivity and for their resting state functional connectivity. An independent t-test between SSD and TDC was performed to compare mean correlational distance across groups for both resting state and EA task mean correlational distance. Additionally, a paired sample t-test between each participant’s mean correlational distance values was run to determine if participants displayed different individual variability during the EA task when compared to the same participants at rest.

### 2.9 Hierarchical Regression Analysis

Hierarchical regression analysis was conducted separately for resting state and EA task individual variability. Hierarchical regression analyses were conducted to examine predictors of individual variability for resting state connectivity and EA background connectivity. Six blocks of independent variables were entered subsequently into seven regression models. We conducted an analysis involving two sets of models with varying orders for entering diagnostic information. Initially, we employed a hierarchical regression where diagnosis was included early in the model with minimal covariates. This aimed to assess the significance of diagnosis while minimizing the influence of other variables. Subsequently, we explored models in which diagnosis was introduced after cognitive and functional outcome variables. This was done to evaluate whether any residual effects of diagnosis remained once cognition was accounted for. The original model included the covariates of age, sex, average FD, and MRI scanner. The second model examined diagnosis, and the third model included a mentalizing factor score (Oliver et al. 2019). Model four added emotional empathy (the IRI empathic concern), model five added the MCCB composite score (Nuechterlein et al. 2008; Keefe et al. 2011), and model six examined functional outcome via subscores of the BSFS. The final model seven added interaction terms, age*sex, age*diagnosis, diagnosis*mentalizing, diagnosis*sex, and mentalizing*sex. We then repeated these hierarchical regressions, entering diagnosis after BSFS scores, but prior to interactions. Examining the change in adjusted-R^2^ statistics between models depicted the amount of variance contributed between steps. ANOVAs were run between each model to determine if the added factors explained significantly more variance in the dependent variable (mean correlational distance).

Two additional hierarchical regression analyses were conducted using only the SSD sample (n = 193) to examine the effects of antipsychotics (CPZ equivalents; (Leucht et al. 2016)), negative symptoms (SANS) and positive symptoms (BPRS) on individual variability, in addition to the independent variables considered in the two previous hierarchical regressions. All independent variables were entered subsequently into ten regression models that were corresponding to ten blocks of independent variables.

### 2.10 Code Sharing

Code used in the analysis of this dataset has been made available (https://github.com/tsecara/Predictors_of_Individual_Variability_SSD)

## 3. Results

### 3.1 Participant Inclusion and Characteristics

Data from 193 individuals with SSD and 155 TDC (total n = 348) were included from the SPINS study (Figure 2). Participant demographics and clinical characteristics, as well as social cognitive factor scores, are shown in Table 1.

**Figure 2.**
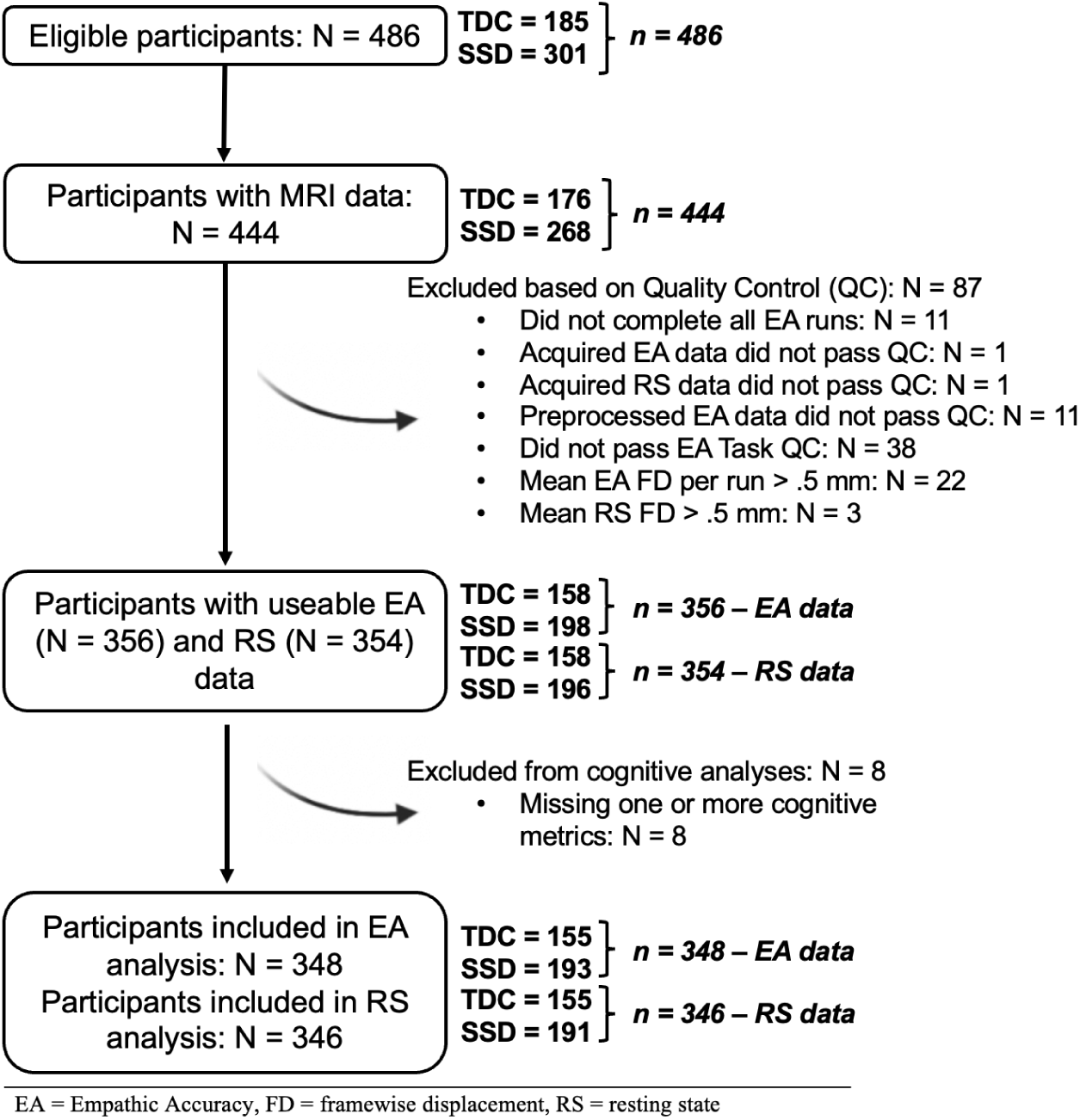
Consort Flow Diagram. Participant data was evaluated for eligibility based on a quality-control (QC) criteria. The initial data from 180 TDC and 286 SSD (n = 466) participants were assessed for available MRI data; 25 participants were excluded. Following, data was controlled for excessive motion and resting state and EA task QC criteria; 87 participants were excluded. Lastly, data was assessed for cognitive score availability; 8 participants were excluded. The resulting data from 155 TDC, 193 SSD individuals EA task data, and 191 SSD individuals with RS data were included for analysis.

**Table 1.** Participant Demographic and Clinical Characteristics by Diagnostic Group.

| Group | SSD<br>(n=193) | TDC<br>(n=155) | P-value |
| --- | --- | --- | --- |
| **Age (years)** | 31.4 (9.6)<br>[18.0, 55.0] | 32.4 (10.6)<br>[18.0, 55.0] | 0.369 |
| **Sex** | 131M (67.9%) | 82M (52.9%) | 0.00618 ** |
| **Education (years)** | 13.8 (2.1) | 15.9 (2.0) | < 2.2e-16 *** |
| **Motion (FD)** | 0.169 (0.094) | 0.138 (0.067) | 0.000361*** |
| **WTAR (standard score)** | 107.30 (14.31) | 113.50 (11.20) | 7.53e-06 *** |
| **BSFS Score** |  |  |  |
| Social engagement/withdrawal | 10.63 (2.33) | 12.50 (1.85) | 1.877e-15 *** |
| Interpersonal | 19.19 (5.35) | 24.78 (3.00) | < 2.2e-16 *** |
| Independence – compliance | 18.09 (10.46) | 32.89 (10.49) | < 2.2e-16 *** |
| Independence – performance | 20.08 (6.58) | 25.55 (7.11) | 1.426e-12 *** |
| Recreation | 28.26 (5.20) | 32.35 (4.19) | 8.575e-15 *** |
| Pro-social | 36.16 (3.14) | 38.35 (1.32) | < 2.2e-16 *** |
| **BPRS Score** |  |  |  |
| Positive symptoms | 7.86 (3.93) | - | - |
| Anxiety and depression | 7.74 (3.53) | - | - |
| Total Score | 31.29 (7.93) | - | - |
| **SANS Score** |  |  |  |
| Affective Flattening | 8.44 (6.02) | - | - |
| Alogia | 1.45 (1.83) | - | - |
| Avolition apathy | 8.57 (5.08) | - | - |
| Anhedonia asociality | 6.70 (4.21) | - | - |
| Total Score | 25.08 (12.55) | - | - |
| **CPZ Equivalents (mg/day)** | 459.10 (397.29) | - | - |
| **MCCB Score** |  |  |  |
| Processing Speed | 40.49 (12.70) | 53.37 (12.71) | < 2.2e-16 *** |
| Attention Vigilance | 39.97 (11.11) | 48.16 (12.14) | 3.214e-10 *** |
| Working Memory | 41.84 (11.18) | 49.61 (11.33) | 5.515e-10 *** |
| Verbal Learning | 41.36 (8.95) | 50.68 (9.23) | < 2.2e-16 *** |
| Visual Learning | 39.98 (12.05) | 48.95 (9.53) | 1.046e-13 *** |
| Reasoning and Problem Solving | 43.83 (10.94) | 48.98 (9.33) | 3.211e-06 *** |
| Total Score | 36.59 (12.19) | 50.12 (10.91) | < 2.2e-16 *** |
| **Scog IRI – Empathic Concern Score** | 20.64 (4.96) | 21.37 (3.86) | 0.126 |
| **Lower-Level Social Cognitive Score (simulation)** | -0.342 (0.86) | 0.378 (0.65) | < 2.2e-16 *** |
| **Higher-Level Social Cognitive Score (mentalizing)** | -0.381 (0.88) | 0.462 (0.62) | < 2.2e-16 *** |

*Note.* Where appropriate variables are displayed as Mean (Standard Deviation). Range presented in square brackets. WTAR= Wechsler Test of Adult Reading, BSFS= Birchwood Social Functioning Score, BPRS= Brief Psychiatric Rating Scale, SANS= Scale for the Assessment of Negative Symptoms, CPZ= Chlorpromazine, MCCB= MATRICS Consensus Cognitive Battery, Scog IRI = Interpersonal Reactivity Index, FD = framewise displacement (\*p < 0.05, \*\*p < 0.01, \*\*\*p < 0.001)

### 3.2 Variability in Network Connectivity

In order to visualise the spatial pattern of regions retained in the analysis, connectivity strength from each region for all retained connections was plotted on the cortical surface parcellated regions, separately for positive and negative connections (Figure 2). The twelve Cole-Anticevic resting-state networks (Ji et al. 2019) were used to identify networks most contributing to the included data. For the EA task, the networks contributing the most to positive background connectivity variability are the default mode, frontoparietal, cingulo-opercular and visual networks (Fig. 2A). For resting state, the visual, default mode and frontoparietal networks display the most variable positive functional connectivity (Fig. 2B). Both the EA task and resting state show similar variability in negative connectivity, with regions overlapping with the cingulo-opercular and default mode being the most implicated (Fig. 2A & Fig. 2B).

**Figure 2.**
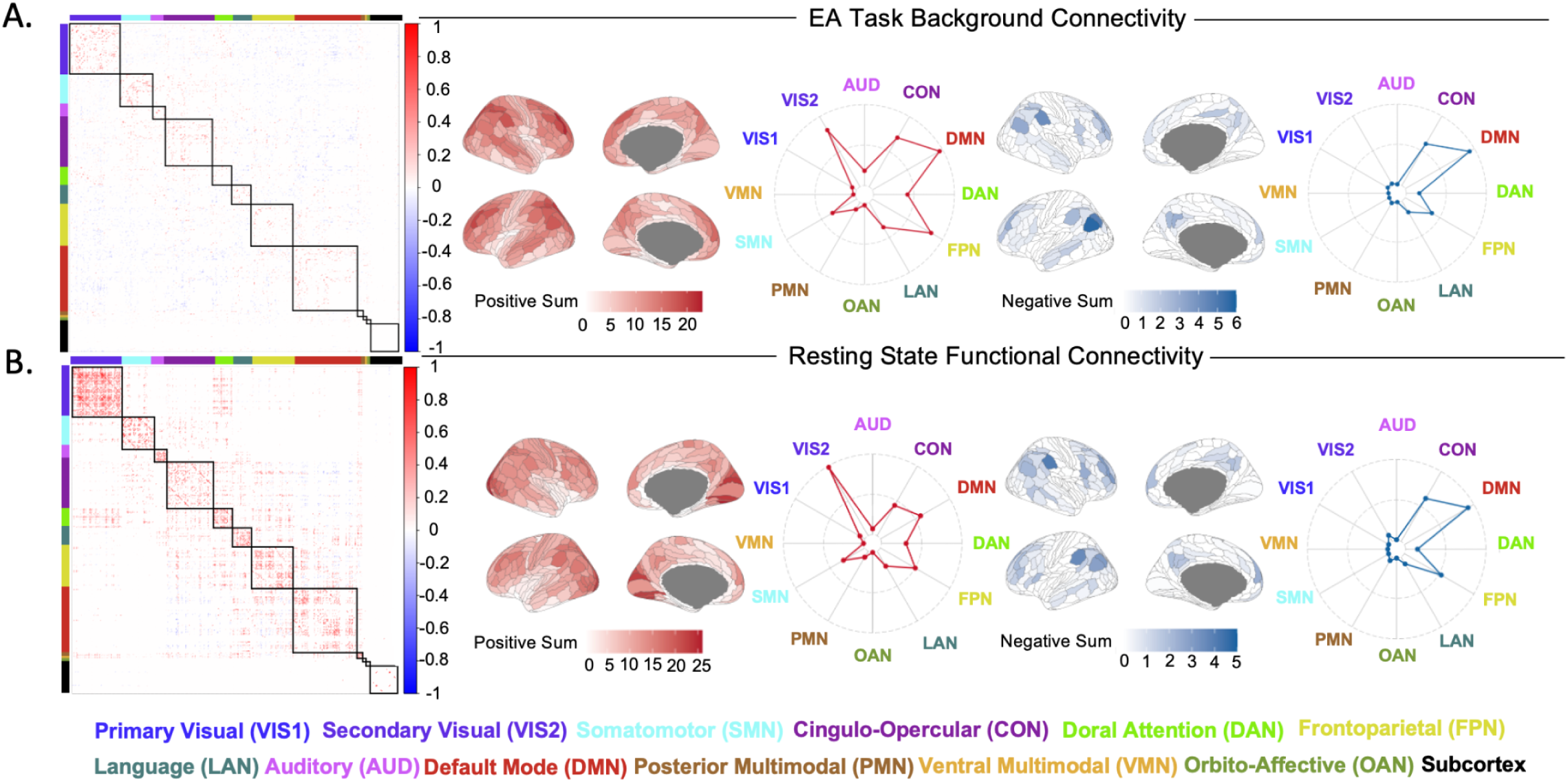
Most Variable Regions of Connectivity: **A)** Background connectivity matrix depicting the top 10% of most variable connections for EA task. Brain visualisations and radar plots depicting the connectivity strength of each network with other retained brain regions, displaying contributions to positive and negative connectivity. **B)** Functional connectivity matrix depicting the top 10% of most variable connections during resting state. Brain visualisations and radar plots depicting the connectivity strength of each network with other retained brain regions, displaying contributions to positive and negative connectivity.

### 3.3 Resting State and EA Task Correlational Distance

Individual correlational distances are plotted in Fig. 3A & 3B. Mean correlational distance of EA task background brain connectivity across SSD (0.5060 ± 0.0042) was found to be significantly higher than TDC (0.5026 ± 0.0040) (Fig. 3C; t = 2.92, p = 0.022, Cohen’s d = 0.24). This was replicated for the resting state data, with mean correlational distance of brain connectivity across SSD (0.5483 ± 0.0019) being significantly higher than TDC (0.5431 ± 0.0018) (Fig. 3D; t = 3.87, p = 0.00013, Cohen’s d = 0.41). Plotting the distribution of mean correlational distance (Fig. 3C & 3D) suggests that SSD and TDC share a substantially overlapping range of mean correlational distance. Mean correlational distance was also significantly lower in EA than in rest (paired-samples t-test; t = 54.65, p = <2.2e-16), demonstrating participants entered into a more similar connectivity pattern while performing the EA task.

**Figure 3.**
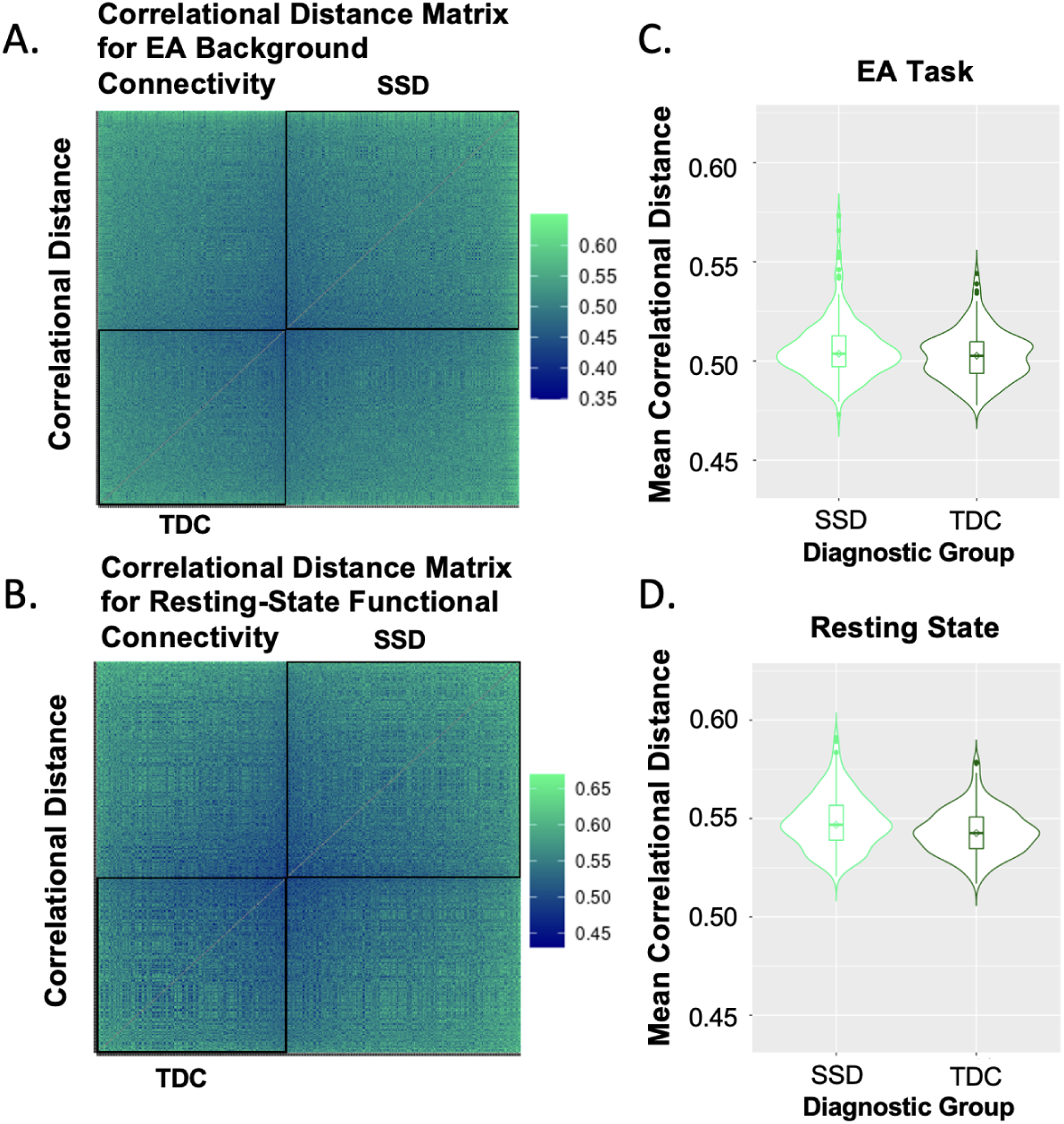
Correlational distance across participants: **A&B)** Pairwise correlational distance between participant pairs was calculated (each row/column represents a single participant, with TDC on the top/left and SSD on the bottom/right). Participants were ordered by mean distance, with TDC in descending and SSD in ascending order, positioning those with the lowest mean distance in the centre of the graph. **C&D)** Violin plots demonstrate between-group differences in mean correlational distance.

### 3.4. Hierarchical Regression Analysis

#### 3.4.1 EA Task

Model 1 included age, sex, average FD, and scanner. These predictors collectively account for 11.58% (adjusted R^2^ = 0.1158) of the variance observed in mean correlational distance (Table 2). Model 2 (diagnostic group) was significant, accounting for an additional 0.95% of variance (adjusted R^2^ = 0.1253, p = 0.031), and Model 3 (mentalizing score) significantly explains an additional 1.44% (adjusted R^2^ = 0.1397, p = 0.010) variance. Model 3 demonstrated that mentalizing (p = 0.016) better accounts for the variability; diagnosis was no longer significant (p = 0.50) when mentalizing was added to the model. Model 4 (IRI empathic concern) is significant and accounts for an additional 1.16% of variance (adjusted R^2^ = 0.1513, p = 0.019). Model 5 (MCCB composite score) and Model 6 (functional outcome scores) did not account for any additional statistically significant variance and was therefore not significant. Model 7 includes interaction terms which account for an additional 1.30% (adjusted R^2^ = 0.1661, p = 0.075), and the final model explains 16.61% of variance in mean correlational distance. Similar results were found in the alternative model with diagnosis added later, after BFSF (Supplemental Table S2). In the final model (Table 2 & Supplemental Table S3), age (F(1, 324) = 10.29, β = 0.00016, p = 0.0015), average framewise displacement (F(1, 324) = 5.12, β = 0.022, p = 0.024), IRI empathic concern (F(1, 324) =3.94, β =0.00033, p = 0.048) and an age by sex interaction term (Fig.4C; F(1, 324) = 4.31, β = −0.00031, p = 0.039) are the only predictor variables that remain significant.

**Figure 4.**
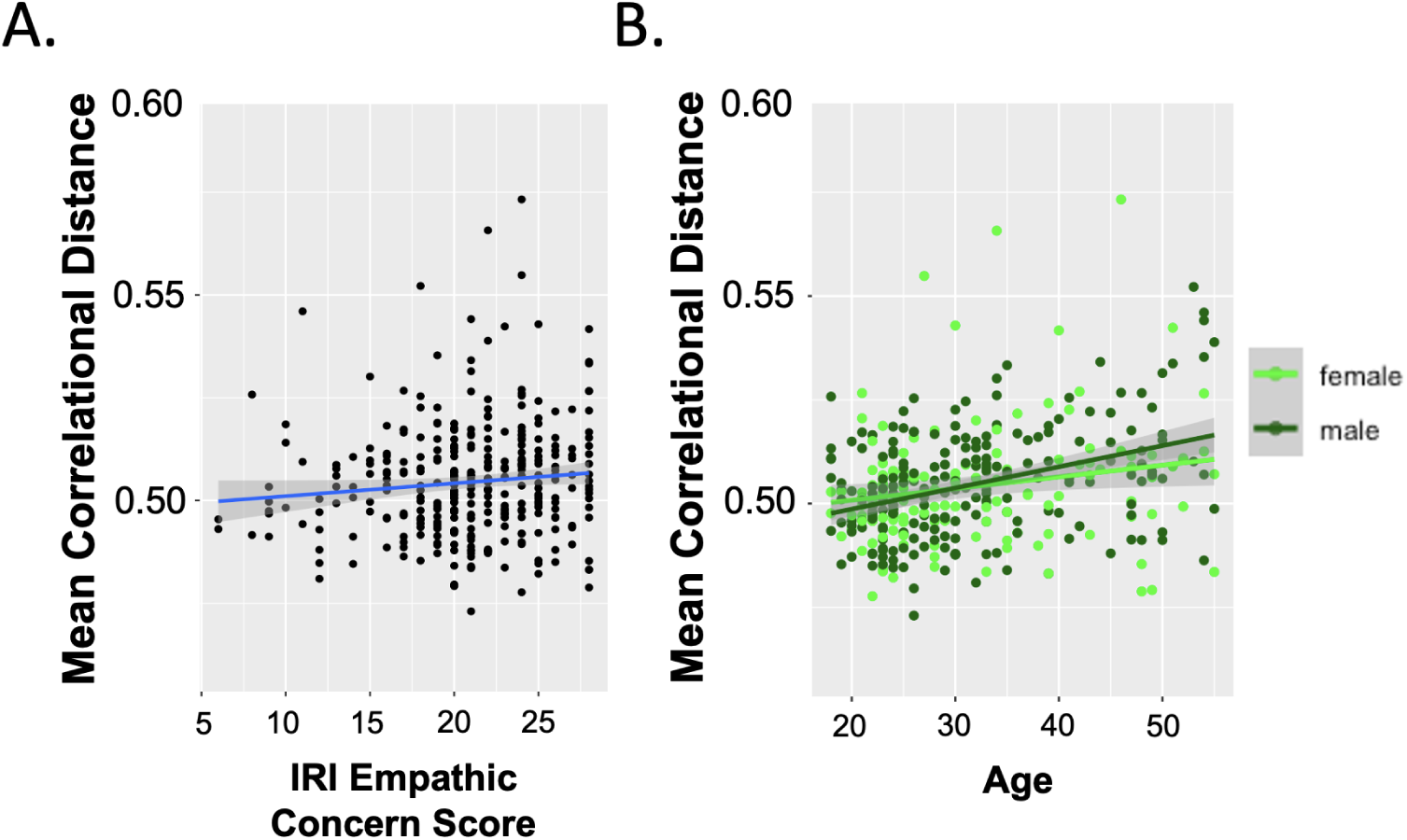
Correlation between predictor variables and EA mean correlational distance: Significant predictor variables and mean correlational distance. Dots represent individual data points, and the darker grey ribbon indicates 95% confidence interval. **A)** Higher mean correlational distance was associated with higher IRI Empathic Concern score. **B)** Increased age was significantly associated with higher mean correlational distance, and this effect was more pronounced in males.

**Table 2:**
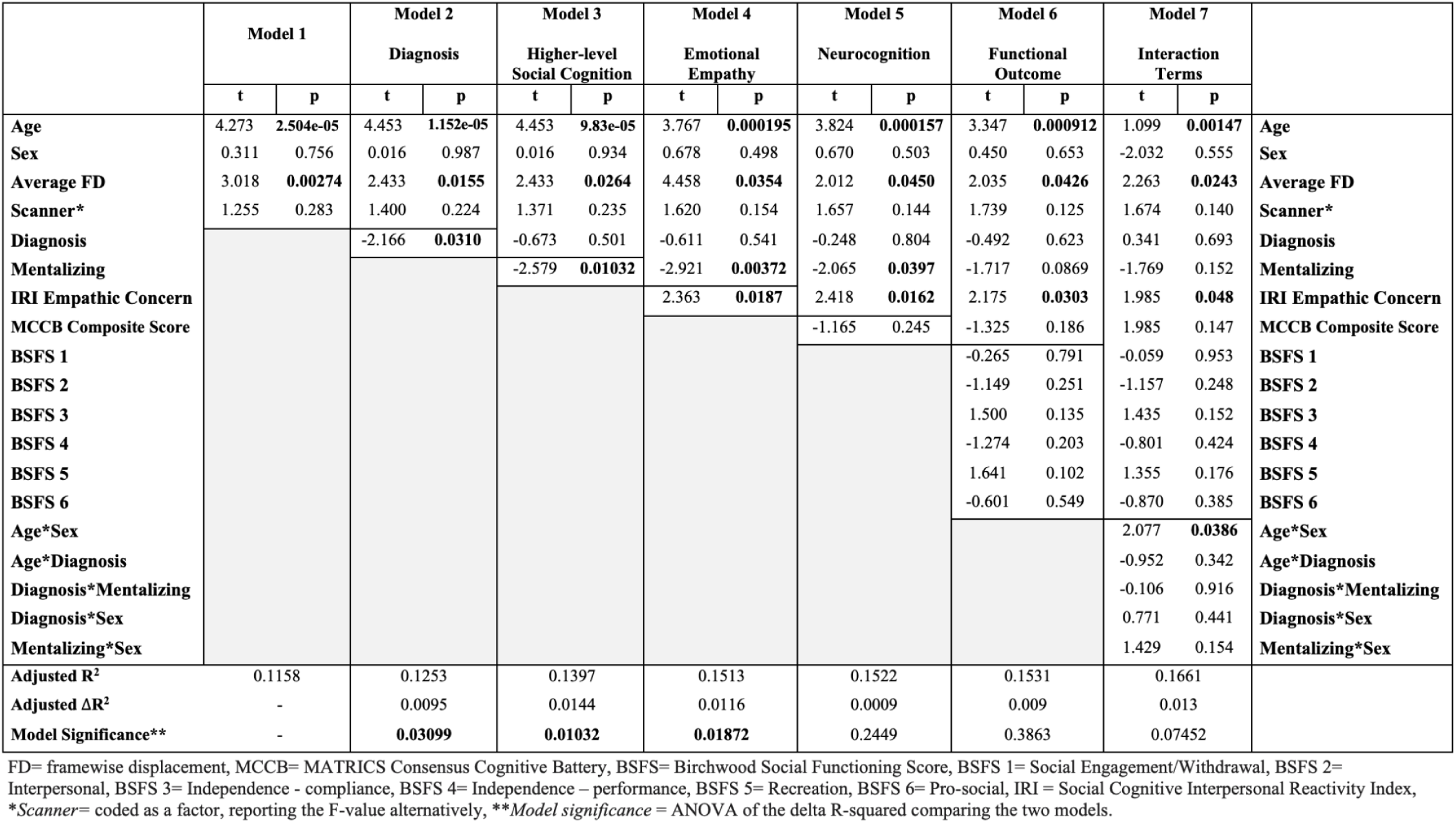
Hierarchical Regression Analysis for EA Task.

#### 3.4.2 Resting State Data

Model 1 (age, sex, average FD, and scanner) accounted for 8.37% (adjusted R^2^ = 0.8367) of the variance observed in mean correlational distance (Table 3). Model 2 (diagnostic group) was statistically significant and accounted for an additional 4.94% of variance (adjusted R^2^ = 0.1331, p = 0.0000096). Model 3 (mentalizing score) was significant as well (adjusted R^2^ = 0.1467, p = 0.012). In contrast to the EA task, diagnosis was still significant in Model 3 (p=0.008), suggesting that diagnostic differences in variability at rest were not accounted for by mentalizing. Model 4 (IRI empathic concern) did not account for any additional statistically significant variance. Model 5 (MCCB composite scores) accounted for an additional 1.17% of variance (adjusted R^2^ = 0.1569, p = 0.018). Model 6 (functional outcome scores) and Model 7 (including interaction terms), both do not significantly account for additional variance.

**Table 3.**
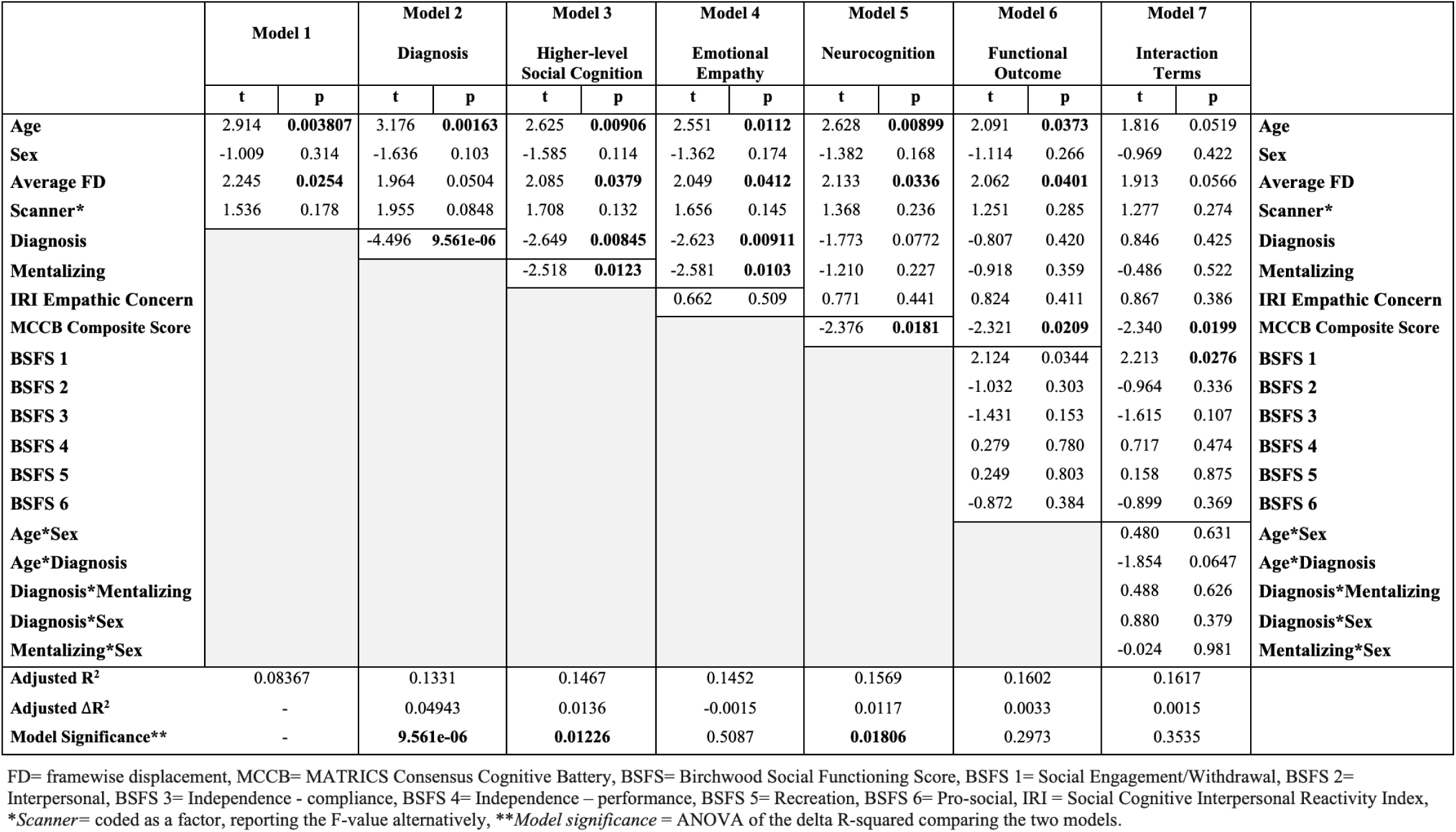
Hierarchical Regression Analysis for Resting State Data.

Interestingly, when diagnosis was entered after cognition and BFSF, mentalizing and MCCB composite score both explained significant additional variance while diagnosis was no longer significant in the hierarchical model (Supplemental Table S4; p = 0.000014 for mentalizing, p = 0.0023 for MCCB composite, and p = 0.42). This suggests similar variance is captured by diagnosis and cognition. The final model explains 16.17% of variance in mean correlational distance. In the final model (Table S5), the BSFS Social Engagement/Withdrawal (F(1, 322) = 5.47, Fig. 5A; β = 0.00070, p = 0.28) and MCCB composite score (F(1, 322) = 4.90, Fig. 5B; β = 0.00077, p = 0.020) were the only predictor variables that remained significant.

**Figure 5.**
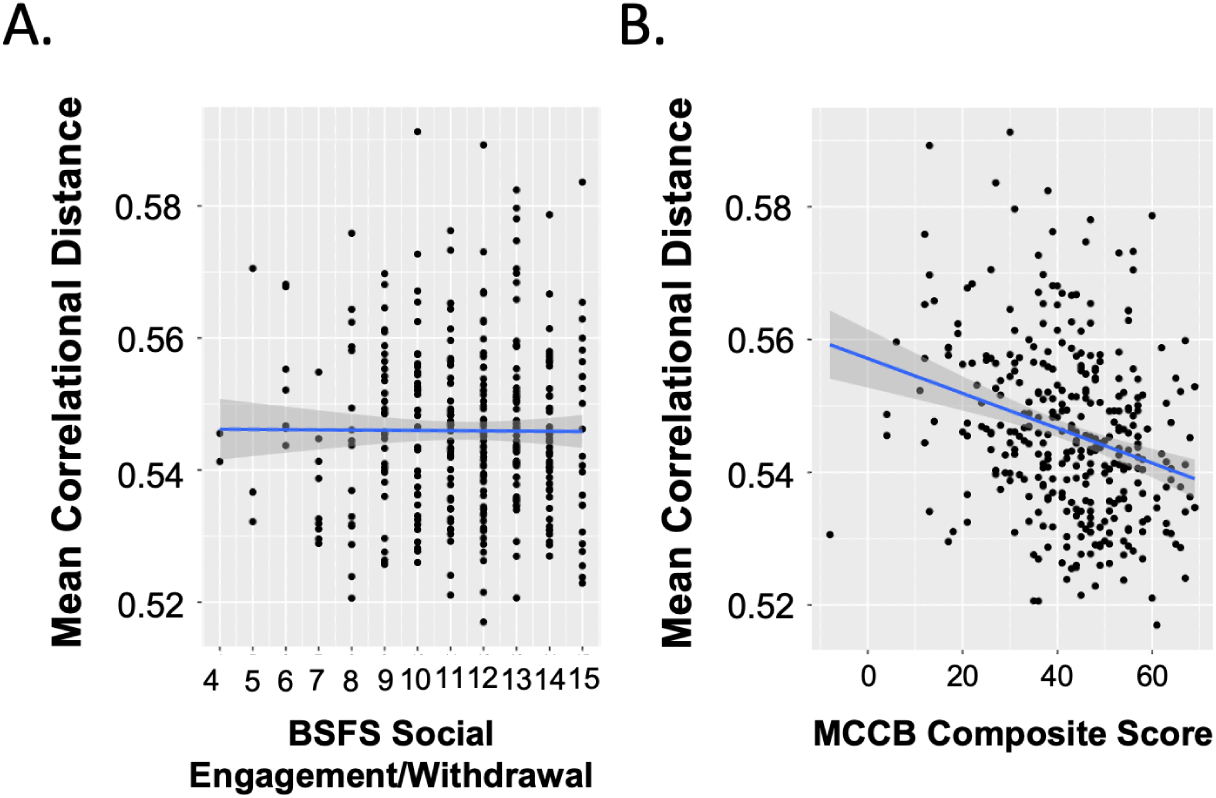
Correlation between predictor variables and resting state mean correlational distance: Significant predictor variables and mean correlational distance. Dots represent individual data points, and the darker grey background indicates 95% confidence interval. **A)** Higher mean correlational distance was significantly associated with higher BSFS Social Engagement/Withdrawal score. **B)** Higher mean correlational distance was associated with lower MCCB composite score.

#### 3.4.3 Clinical Scores and Medication Effects in the SSD group

When examining the effects of medication (CPZ equivalents) and clinical scores (SANS negative symptoms, and BPRS positive symptoms and depression/anxiety) in the SSD group, medication effects and clinical scores did not contribute additional variance (See Supplementary tables S6 and S7).

## 4. Discussion

In the present study, we investigated predictors of individual variability in task and resting functional connectivity in a large sample of SSD and TDC. Individuals displayed greater average variability during resting state compared to the EA task; the observed increase in individual variability may be attributed to the unconstrained nature of resting state, where participants engage in mind wandering and spontaneous, internally directed cognitive processes (Diaz et al. 2013; Marchetti et al. 2015), whereas the task may produce a more constrained state (Cole et al. 2014). While initial comparisons revealed diagnostic differences in mean correlational distance between SSD and TDC during the EA task, hierarchical regression analysis revealed that variability in EA task background connectivity was related to mentalizing score and empathy, but not diagnostic group, while resting state variability was predominantly related to diagnostic differences. Lower cognitive scores were related to increased variability, suggesting that poor performers exhibit an underlying pattern of globally unstable connectivity or dysconnectivity (Cole et al. 2011). These results suggest that unconstrained resting fMRI may emphasize diagnostic differences in SSD, while cognitive tasks map onto the relevant behaviors.

Studying individual variability during resting state emphasizes diagnostic differences in functional connectivity between SSD and TDC. Several hypotheses have suggested dysconnectivity in SSD due to abnormal functional integration (K. J. Friston and Frith 1995; Weinberger 1993; K. Friston et al. 2016), or global dysconnectivity linked to symptomology in SSD (Cole et al. 2011). The dysconnectivity hypothesis may extend to the internally directed cognitive processes and self-referential nature of resting state, directly translating to the disorganized and increased variability observed in the SSD sample. Individuals with SSD display greater variability during resting state compared to TDC, while during the EA task diagnostic effects were not present once mentalizing was included in the model. This suggests that resting state is an optimal modality for studying diagnostic differences and a powerful tool for observing the consistent abnormalities in the functional organization and variability that characterizes SSD (Sheffield and Barch 2016; Woodward, Rogers, and Heckers 2011).

While group differences in EA task variability between SSD and TDC were found, aligning with recent work examining variability in task fMRI showing greater variability in SSD (Hawco et al. 2020; Gallucci, Tan, et al. 2022; Gallucci, Pomarol-Clotet, et al. 2022), no significant group differences were observed when social cognition was accounted for in the EA task analysis. The current results fall in line with our recent work suggesting that social cognitive functional task-activity or connectivity in SSD falls along a similar spectrum of connectivity as controls (Oliver et al. 2021; Viviano et al. 2018; Hawco et al. 2019), with SSD more likely to be on the ‘impaired’ end of the spectrum. When performing group comparisons across SSD and TDC, factors such as cognition may represent confounds given that cognitive differences in neurobiology can masquerade as diagnostic differences if not accounted for in the analyses. The EA task engages brain regions within the mentalizing network (Oliver et al. 2021; Zaki et al. 2009; Harvey et al. 2013), which shows overlap with the default mode network (Alcalá-López et al. 2019). Differences in mentalizing were significantly related to variability differences during the EA task. The EA task shifts connectivity into more constrained social cognitive brain states elicited by neural strategies underlying emotional-processing (Cole et al. 2014; Oliver et al. 2021). Original work by (Zaki et al. 2009) suggests that the EA task evokes activity in multiple networks, proposing that parallel activation of both mirror neuron and mental state attribution systems correspond to EA. Poorer social cognitive performers may show social network impairments, necessitating the use of alternate pathways to complete the task. Alternatively, poor performers may use less efficient task strategies during the task performance, corresponding to different neural networks, resulting in greater variability (Oliver et al. 2021; Kanske et al. 2016; Hawco et al. 2019).

Several limitations of our study should be noted. Motion continues to be a challenge in many studies of brain structure and function (Savalia et al. 2017; Van Dijk, Sabuncu, and Buckner 2012). Our study aimed to account for this confound by correcting for motion at several stages of the analysis. Furthermore, there are additional potential sources of variability that were not examined here, which include but are not limited to: body mass index (Kullmann et al. 2012), genetic variants (Gilman et al. 2012), fluid and crystalized intelligence (Kruschwitz et al. 2018), personality domains (Hsu et al. 2018), sleep quality (Sämann et al. 2010), and illness duration (Gallucci, Pomarol-Clotet, et al. 2022). This may account for the large proportion of variance that remains unexplained in the resting state and EA task hierarchical regression.

To our knowledge, this study was the first to use mean correlational distance as a global metric of individual variability during resting state and social cognitive task-based fMRI in SSD. Apparent diagnostic differences in functional connectivity variability in SSD and TDC were driven by differences in cognitive abilities during the EA task, in line with previous findings by our group (Oliver et al. 2021), while resting state strongly elucidated diagnostic differences in variability. Examining connectivity during a social cognitive processing task state allowed for better delineation of the relationship between variability and higher level social cognitive performance, while resting state emphasized diagnostic differences related to variability. Future work should consider both resting state and task-state connectivity when examining variability. Further validation of the variability underlying brain-behavior relationships could guide targeted treatment development for those exhibiting cognitive deficits, and future studies should consider individualized patterns of variability during task and rest to optimize individualized treatment.

