## Supplementary Information for "Heterogeneity in Functional Connectivity: Dimensional Predictors of Individual Variability during Rest and Task fMRI in Psychosis"

##### Supplementary Methods and Materials

**Empathic Accuracy (EA) Task**
The EA task was completed during functional MRI. Participants were trained using a practice version of the task in a mock scanner prior to scanning. During the task, participants watch nine videos discussing emotional (4 positive, 5 negative) autobiographical events. This version of the EA task was designed to include adults varying in age, race, and ethnicity, and details regarding video development are presented elsewhere (1, 2). EA was calculated for each participant by correlating their ratings with self-ratings provided by individuals in the videos, and EA values were then Fisher r-to-z transformed. During the control condition, which was two interleaved control videos (40 s each) per run, participants provide continuous ratings of the relative light or darkness of a grayscale circle as it changes shades, on a 9-point scale. This condition is included to ensure that participants are engaged in the task (controlling for avolition) and comprehend it.

**MRI Data Acquisition**
MRI scans were collected using harmonized scanning parameters on 3T scanners with multichannel head coils, including a General Electric Discovery (N=113; CAMH) and Siemens Prisma (N=27; CAMH), a General Electric Signa (N=28; ZHH) and Siemens Prisma (N=66; ZHH), and a Siemens Tim Trio (N=52; MPRC) and Siemens Prisma (N=62; MPRC).

**EA Task GLM Nuisance Regressors**
To model the stimulus-evoked response, each of the nine EA videos were fitted with three hemodynamic response functions (HRF): one for the video duration, another modulated by the participant's EA score, and a third for button presses made during the video (lasting 1 second). Control videos were also included and modeled with two HRFs: one for the video duration and another for button presses made during the video (lasting 1 second). These regressors were applied to each voxel, and the resulting residual activation was used for background connectivity analysis (3).

**MRI Exclusion Criteria**
Prior to analyses, all scans were quality checked by experienced research staff, using an in-house quality control system dashboard (https://github.com/TIGRLab/dashboard) before preprocessing, and after preprocessing (4, 5). This included qualitative (e.g., detection of ghosting or ringing) and quantitative (e.g., framewise displacement) monitoring. Participants were also excluded for excessive motion (mean framewise displacement per run > 0.5 mm) during the EA task or RS scan, and EA or control task performance indicative of disengagement or a lack of comprehension. Specifically, participants were excluded if they had any EA videos with no responses, or more than one video with only one response, or a control task accuracy of < 20%

and one or two responses in any EA video. As mentioned, participants were also excluded from analyses if they were missing any cognitive metrics. See the consort flow diagram for inclusion/exclusion details (Figure 2).

**fMRI Preprocessing**

Anatomical T1-weighted images were corrected for intensity non-uniformity using ANTS 2.2.0 (6) and skull-stripping using Nipype implementation of the antsBrainExtraction.sh workflow (from ANTs). Brain tissue segmentation of cerebrospinal fluid, white-matter and gray-matter was performed using FSL 5.0.9 (7). Brain surfaces were reconstructed using FreeSurfer 6.0.1 (8). The extracted brain was spatially normalized to MNI space using nonlinear registration with ANTs (6). For each of the fMRI runs, fieldmap-less distortion correction was performed using fMRIPrep. Registration is performed using ANTs (9). Functional data was coregistered to the corresponding T1-weighted image using Freesurfer's boundary-based registration with six degrees of freedom. Head-motion parameters were estimated using MCFLIRT (FSL 5.0.9; (10)), and slice-time correction was performed using 3dTshift from AFNI (11), RRID_SCR_005927). The Ciftify toolbox (12), https://github.com/edickie/ciftify) was used to transform the functional data onto the cortical surface (fsLR32k space; (13)) using a non-linear transform to the MNI152 template via FSL's FNIRT. Four TRs were dropped for each EA scan and three TRs for the resting state scan, and data was smoothed at 2mm FWHM on the cortical surface. For resting state, detrending, band-pass filtering (0.01-0.1 Hz), and nuisance regression was also performed using Ciftify. Detrending and nuisance regression occurred within the GLM for the EA task data. For both resting state and EA data, the nuisance regression model included regressors for the six head motion correction parameters, mean white matter signal, mean cerebral spinal fluid signal, the square, derivative, and square of the derivative for each of these regressors, and global brain signal (generated by fMRIPrep) (14, 15). For the EA task, an amplitude-modulated general linear model was performed using AFNI's 3dDeconvolve module in nistats (16, 11). To model the stimulus-evoked response, the regressors were fit to each voxel and the residual activation was retained for background connectivity analysis (17, 3). Background connectivity allows an examination of state-related connectivity rather than stimulus-driven coactivation (17, 18).

### Supplementary Results

#### Table S1: Correlation Matrix of Variables Used in Hierarchical Regression Analyses

| | Site | Scanner | Diagnostic Group | Sex | Age | Mean CorDist EA | Mean CorDist EA* | Mean CorDist RS | Avg FD EA | Avg FD RS | Simulation | Mentalizing | WTAR | Empathic Concern | Processing Speed | Attention Vigilance | Working Memory | Verbal Learning | Visual Learning | Reasoning & PS | BSFS Sec 1 | BSFS Sec 2 | BSFS Sec 3 | BSFS Sec 4 | BSFS Sec 5 | BSFS Sec 6 |
|---|---|---|---|---|---|---|---|---|---|---|---|---|---|---|---|---|---|---|---|---|---|---|---|---|---|---|
| Site | 1 | 0.97 | -0.12 | 0.04 | 0.32 | 0.22 | 0.16 | 0.19 | 0.23 | 0.15 | -0.18 | -0.11 | -0.18 | 0.06 | -0.05 | 0.03 | -0.13 | -0.03 | -0.11 | -0.03 | 0.12 | 0.08 | -0.02 | -0.03 | 0.07 | 0.01 |
| Scanner | 0.97 | 1 | -0.12 | 0.05 | 0.29 | 0.22 | 0.18 | 0.17 | 0.24 | 0.16 | -0.16 | -0.09 | -0.16 | 0.03 | -0.03 | 0.04 | -0.11 | 0 | -0.1 | -0.01 | 0.12 | 0.1 | 0 | -0.02 | 0.07 | -0.01 |
| Diagnostic Group | -0.12 | -0.12 | 1 | -0.15 | -0.06 | 0.14 | 0.12 | 0.2 | 0.17 | 0.06 | -0.42 | -0.47 | -0.23 | -0.07 | -0.49 | -0.33 | -0.32 | -0.45 | -0.38 | -0.25 | -0.4 | -0.53 | -0.58 | -0.37 | -0.39 | -0.4 |
| Sex | 0.04 | 0.05 | -0.15 | 1 | 0.01 | -0.03 | -0.01 | 0.04 | -0.04 | -0.09 | 0.05 | 0.04 | -0.08 | 0.25 | 0.07 | 0.01 | 0.09 | 0.03 | 0.08 | 0.03 | 0.19 | 0.19 | 0.07 | 0.09 | 0.15 | 0.07 |
| Age | 0.32 | 0.29 | -0.06 | 0.01 | 1 | 0.39 | 0.29 | 0.26 | 0.33 | 0.3 | -0.24 | -0.17 | -0.16 | 0.1 | 0.05 | 0.11 | -0.08 | -0.14 | -0.06 | 0.01 | 0.05 | 0.01 | -0.15 | -0.07 | 0.24 | 0.04 |
| Mean CorDist EA | 0.22 | 0.22 | 0.14 | -0.03 | 0.39 | 1 | 0.84 | 0.45 | 0.57 | 0.38 | -0.38 | -0.3 | -0.24 | 0.14 | -0.19 | -0.04 | -0.18 | -0.19 | -0.17 | -0.13 | -0.05 | -0.07 | -0.1 | -0.07 | 0.09 | -0.08 |
| Mean CorDist EA* | 0.16 | 0.18 | 0.12 | -0.01 | 0.29 | 0.84 | 1 | 0.44 | 0.25 | 0.21 | -0.31 | -0.23 | -0.17 | 0.12 | -0.17 | -0.02 | -0.17 | -0.14 | -0.12 | -0.17 | -0.04 | -0.08 | -0.07 | -0.08 | 0.1 | -0.07 |
| Mean CorDist RS | 0.19 | 0.17 | 0.2 | 0.04 | 0.26 | 0.45 | 0.44 | 1 | 0.21 | 0.19 | -0.3 | -0.26 | -0.28 | 0.06 | -0.26 | -0.09 | -0.22 | -0.2 | -0.25 | -0.23 | -0.01 | -0.16 | -0.23 | -0.12 | -0.03 | -0.14 |
| Avg FD EA | 0.23 | 0.24 | 0.17 | -0.04 | 0.33 | 0.57 | 0.25 | 0.21 | 1 | 0.57 | -0.24 | -0.2 | -0.19 | 0.06 | -0.14 | -0.08 | -0.12 | -0.16 | -0.16 | -0.05 | -0.15 | -0.12 | -0.19 | -0.1 | -0.04 | -0.08 |
| Avg FD RS | 0.15 | 0.16 | 0.06 | -0.09 | 0.3 | 0.38 | 0.21 | 0.19 | 0.57 | 1 | -0.09 | -0.04 | -0.07 | 0.04 | 0.04 | 0.06 | -0.01 | -0.04 | -0.06 | 0.1 | -0.1 | -0.05 | -0.08 | -0.02 | 0.05 | -0.17 |
| Simulation | -0.18 | -0.16 | -0.42 | 0.05 | -0.24 | -0.38 | -0.31 | -0.3 | -0.24 | -0.09 | 1 | 0.85 | 0.55 | 0.12 | 0.46 | 0.39 | 0.49 | 0.46 | 0.46 | 0.35 | 0.12 | 0.3 | 0.34 | 0.25 | 0.11 | 0.24 |
| Mentalizing | -0.11 | -0.09 | -0.47 | 0.04 | -0.17 | -0.3 | -0.23 | -0.26 | -0.2 | -0.04 | 0.85 | 1 | 0.5 | 0.11 | 0.46 | 0.38 | 0.48 | 0.48 | 0.44 | 0.36 | 0.18 | 0.35 | 0.37 | 0.3 | 0.15 | 0.27 |
| WTAR | -0.18 | -0.16 | -0.23 | -0.08 | -0.16 | -0.24 | -0.17 | -0.28 | -0.19 | -0.07 | 0.55 | 0.5 | 1 | -0.01 | 0.41 | 0.31 | 0.41 | 0.38 | 0.45 | 0.27 | 0 | 0.16 | 0.18 | 0.17 | 0.13 | 0.19 |
| Empathic Concern | 0.06 | 0.03 | -0.07 | 0.25 | 0.1 | 0.14 | 0.12 | 0.06 | 0.06 | 0.04 | 0.12 | 0.11 | -0.01 | 1 | 0.05 | 0.14 | -0.02 | 0 | 0.06 | -0.01 | 0.16 | 0.18 | 0.12 | 0.06 | 0.17 | 0.04 |
| Processing Speed | -0.05 | -0.03 | -0.49 | 0.07 | 0.05 | -0.19 | -0.17 | -0.26 | -0.14 | 0.04 | 0.46 | 0.46 | 0.41 | 0.05 | 1 | 0.5 | 0.6 | 0.53 | 0.55 | 0.53 | 0.22 | 0.32 | 0.34 | 0.21 | 0.29 | 0.26 |
| Attention Vigilance | 0.03 | 0.04 | -0.33 | 0.01 | 0.11 | -0.04 | -0.02 | -0.09 | -0.08 | 0.06 | 0.39 | 0.38 | 0.31 | 0.14 | 0.5 | 1 | 0.45 | 0.4 | 0.37 | 0.3 | 0.12 | 0.19 | 0.15 | 0.05 | 0.17 | 0.17 |
| Working Memory | -0.13 | -0.11 | -0.32 | 0.09 | -0.08 | -0.18 | -0.17 | -0.22 | -0.12 | -0.01 | 0.49 | 0.48 | 0.41 | -0.02 | 0.6 | 0.45 | 1 | 0.45 | 0.53 | 0.43 | 0.13 | 0.22 | 0.29 | 0.22 | 0.15 | 0.2 |
| Verbal Learning | -0.03 | 0 | -0.45 | 0.03 | -0.14 | -0.19 | -0.14 | -0.2 | -0.16 | -0.04 | 0.46 | 0.48 | 0.38 | 0 | 0.53 | 0.4 | 0.45 | 1 | 0.56 | 0.27 | 0.15 | 0.26 | 0.33 | 0.23 | 0.22 | 0.25 |
| Visual Learning | -0.11 | -0.1 | -0.38 | 0.08 | -0.06 | -0.17 | -0.12 | -0.25 | -0.16 | -0.06 | 0.46 | 0.44 | 0.45 | 0.06 | 0.55 | 0.37 | 0.53 | 0.56 | 1 | 0.32 | 0.14 | 0.2 | 0.24 | 0.22 | 0.18 | 0.24 |
| Reasoning & PS | -0.03 | -0.01 | -0.25 | 0.03 | 0.01 | -0.13 | -0.17 | -0.23 | -0.05 | 0.1 | 0.35 | 0.36 | 0.27 | -0.01 | 0.53 | 0.3 | 0.43 | 0.27 | 0.32 | 1 | 0.07 | 0.2 | 0.18 | 0.19 | 0.16 | 0.19 |
| BSFS Sec 1 | 0.12 | 0.12 | -0.4 | 0.19 | 0.05 | -0.05 | -0.04 | -0.01 | -0.15 | -0.1 | 0.12 | 0.18 | 0 | 0.16 | 0.22 | 0.12 | 0.13 | 0.15 | 0.14 | 0.07 | 1 | 0.53 | 0.43 | 0.32 | 0.4 | 0.3 |
| BSFS Sec 2 | 0.08 | 0.1 | -0.53 | 0.19 | 0.01 | -0.07 | -0.08 | -0.16 | -0.12 | -0.05 | 0.3 | 0.35 | 0.16 | 0.18 | 0.32 | 0.19 | 0.22 | 0.26 | 0.2 | 0.2 | 0.53 | 1 | 0.56 | 0.33 | 0.36 | 0.3 |
| BSFS Sec 3 | -0.02 | 0 | -0.58 | 0.07 | -0.15 | -0.1 | -0.07 | -0.23 | -0.19 | -0.08 | 0.34 | 0.37 | 0.18 | 0.12 | 0.34 | 0.15 | 0.29 | 0.33 | 0.24 | 0.18 | 0.43 | 0.56 | 1 | 0.6 | 0.44 | 0.29 |
| BSFS Sec 4 | -0.03 | -0.02 | -0.37 | 0.09 | -0.07 | -0.07 | -0.08 | -0.12 | -0.1 | -0.02 | 0.25 | 0.3 | 0.17 | 0.06 | 0.21 | 0.05 | 0.22 | 0.23 | 0.22 | 0.19 | 0.32 | 0.33 | 0.6 | 1 | 0.49 | 0.29 |
| BSFS Sec 5 | 0.07 | 0.07 | -0.39 | 0.15 | 0.24 | 0.09 | 0.1 | -0.03 | -0.04 | 0.05 | 0.11 | 0.15 | 0.13 | 0.17 | 0.29 | 0.17 | 0.15 | 0.22 | 0.18 | 0.16 | 0.4 | 0.36 | 0.44 | 0.49 | 1 | 0.32 |
| BSFS Sec 6 | 0.01 | -0.01 | -0.4 | 0.07 | 0.04 | -0.08 | -0.07 | -0.14 | -0.08 | -0.17 | 0.24 | 0.27 | 0.19 | 0.04 | 0.26 | 0.17 | 0.2 | 0.25 | 0.24 | 0.19 | 0.3 | 0.3 | 0.29 | 0.29 | 0.32 | 1 |

EA = Empathic Accuracy, RS = resting state, FD = framewise displacement, CorDist = correlational distance, WTAR= Wechsler Test of Adult Reading, PS = problem solving, BSFS = Birchwood Social Functioning Scale, * = motion reduced

### Table S2: Hierarchical Regression Analysis for EA task including Diagnostic Group as Model 6

| | Model 1 | | Model 2 Higher-level Social Cognition | | Model 3 Emotional Empathy | | Model 4 Neurocognition | | Model 5 Functional Outcome | | Model 6 Diagnosis | | Model 7 Interaction Terms | | |
|---|---|---|---|---|---|---|---|---|---|---|---|---|---|---|---|
| | t | p | t | p | t | p | t | p | t | p | t | p | t | p | |
| **Age** | 4.273 | **2.504e-05** | 3.888 | **0.000122** | 3.720 | **0.000233** | 3.831 | **0.000152** | 3.314 | **0.00102** | 3.347 | **0.000912** | 1.099 | **0.00147** | **Age** |
| **Sex** | 0.311 | 0.756 | 0.173 | 0.863 | 0.767 | 0.444 | 0.708 | 0.479 | 0.499 | 0.618 | 0.450 | 0.653 | -2.032 | 0.555 | **Sex** |
| **Average FD** | 3.018 | **0.00274** | 2.378 | **0.0180** | 2.245 | **0.0254** | 2.067 | **0.0395** | 2.086 | **0.0377** | 2.035 | **0.0426** | 2.263 | **0.0243** | **Average FD** |
| **Scanner** | 1.255 | 0.283 | 1.337 | 0.248 | 1.593 | 0.161 | 1.654 | 0.145 | 1.753 | 0.122 | 1.738 | 0.125 | 1.674 | 0.140 | **Scanner** |
| **Mentalizing** | | | -3.315 | **0.00102** | -3.668 | **0.000284** | -2.222 | **0.0270** | -1.842 | 0.0663 | -1.717 | 0.0869 | -1.769 | 0.152 | **Mentalizing** |
| **IRI Empathic Concern** | | | | | 2.383 | **0.0177** | 2.433 | **0.0155** | 2.251 | **0.0250** | 2.175 | **0.0303** | 1.985 | **0.0480** | **IRI Empathic Concern** |
| **MCCB Composite Score** | | | | | | | -1.294 | 0.197 | -1.464 | 0.144 | -1.325 | 0.186 | -1.455 | 0.147 | **MCCB Composite Score** |
| **BSFS 1** | | | | | | | | | -0.290 | 0.772 | -0.265 | 0.791 | -0.059 | 0.953 | **BSFS 1** |
| **BSFS 2** | | | | | | | | | -1.239 | 0.216 | -1.149 | 0.251 | -1.157 | 0.248 | **BSFS 2** |
| **BSFS 3** | | | | | | | | | 1.419 | 0.157 | 1.500 | 0.135 | 1.435 | 0.152 | **BSFS 3** |
| **BSFS 4** | | | | | | | | | -1.255 | 0.210 | -1.274 | 0.203 | -0.801 | 0.424 | **BSFS 4** |
| **BSFS 5** | | | | | | | | | 1.625 | 0.105 | 1.641 | 0.102 | 1.355 | 0.176 | **BSFS 5** |
| **BSFS 6** | | | | | | | | | -0.696 | 0.487 | -0.601 | 0.549 | -0.870 | 0.385 | **BSFS 6** |
| **Diagnosis** | | | | | | | | | | | -0.492 | 0.623 | 0.341 | 0.693 | **Diagnosis** |
| **Age*Sex** | | | | | | | | | | | | | 2.077 | **0.0386** | **Age*Sex** |
| **Age*Diagnosis** | | | | | | | | | | | | | -0.952 | 0.342 | **Age*Diagnosis** |
| **Diagnosis*Mentalizing** | | | | | | | | | | | | | -0.106 | 0.916 | **Diagnosis*Mentalizing** |
| **Diagnosis*Sex** | | | | | | | | | | | | | 0.771 | 0.441 | **Diagnosis*Sex** |
| **Mentalizing*Sex** | | | | | | | | | | | | | 1.429 | 0.154 | **Mentalizing*Sex** |
| **Adjusted $R^2$** | 0.1158 | | 0.1411 | | 0.1528 | | 0.1545 | | 0.1550 | | 0.1531 | | 0.1661 | | |
| **Adjusted $\Delta R^2$** | - | | 0.0253 | | 0.0117 | | 0.0017 | | 0.0005 | | -0.0019 | | 0.013 | | |
| **Model Significance** | - | | **0.001016** | | **0.01773** | | 0.1966 | | 0.4038 | | 0.6234 | | 0.07452 | | |

FD= framewise displacement, MCCB= MATRICS Consensus Cognitive Battery, BSFS= Birchwood Social Functioning Score, BSFS 1= Social Engagement/Withdrawal, BSFS 2= Interpersonal, BSFS 3= Independence - compliance, BSFS 4= Independence – performance, BSFS 5= Recreation, BSFS 6= Pro-social, IRI = Social Cognitive Interpersonal Reactivity Index, **Scanner*= coded as a factor, reporting the F-value alternatively, ***Model significance* = ANOVA of the delta R-squared comparing the two models.

#### Table S3: Significant Predictor Variables in Final Model of EA Task Hierarchical Regression

| Predictor Variable | β coefficient | F-value | P-value |
|---|---|---|---|
| **Age** | 0.0001648 | 10.2946 | **0.00147** |
| Sex | -0.01039 | 0.3499 | 0.555 |
| **Average FD** | 0.02158 | 5.1218 | **0.0243** |
| Scanner | -0.0002368 | 1.6740 | 0.140 |
| Diagnostic group | 0.001836 | 0.1560 | 0.693 |
| Mentalizing | -0.003546 | 2.0573 | 0.152 |
| **IRI Empathic Concern** | 0.0003284 | 3.9385 | **0.048** |
| MCCB Composite Score | -0.0001012 | 2.1178 | 0.147 |
| BSFS Social Engagement/Withdrawal | -0.00002223 | 0.0035 | 0.953 |
| BSFS Interpersonal | -0.0002118 | 1.3387 | 0.248 |
| BSFS Independence-compliance | 0.0001222 | 2.0601 | 0.152 |
| BSFS Independence-performance | -0.0001031 | 0.6409 | 0.424 |
| BSFS Recreation | 0.0002416 | 1.8360 | 0.176 |
| BSFS Pro-social | -0.0002567 | 0.7572 | 0.385 |
| **Age*sex** | 0.0003077 | 4.3148 | **0.0386** |
| Age*diagnostic group | -0.0001393 | 0.9070 | 0.342 |
| Diagnostic group*mentalizing | -0.0002226 | 0.0112 | 0.916 |
| Diagnostic group*sex | 0.002674 | 0.5948 | 0.441 |
| Mentalizing*sex | 0.003032 | 2.0419 | 0.154 |

FD= framewise displacement, MCCB= MATRICS Consensus Cognitive Battery, BSFS= Birchwood Social Functioning Score, IRI = Social Cognitive Interpersonal Reactivity Index

### Table S4: Hierarchical Regression Analysis for RS including Diagnostic Group as Model 6

| | Model 1 | | Model 2 Higher-level Social Cognition | | Model 3 Emotional Empathy | | Model 4 Neurocognition | | Model 5 Functional Outcome | | Model 6 Diagnosis | | Model 7 Interaction Terms | | |
|---|---|---|---|---|---|---|---|---|---|---|---|---|---|---|---|
| | t | p | t | p | t | p | t | p | t | p | t | p | t | p | |
| **Age** | 2.914 | **0.00381** | 2.265 | **0.0242** | 2.189 | **0.0293** | 2.419 | **0.0161** | 2.004 | **0.0459** | 2.091 | **0.0373** | 1.816 | 0.0519 | **Age** |
| **Sex** | -1.009 | 0.314 | -1.240 | 0.216 | -1.011 | 0.313 | -1.164 | 0.245 | -1.042 | 0.298 | -1.114 | 0.266 | -0.969 | 0.422 | **Sex** |
| **Average FD** | 2.245 | **0.0254** | 2.301 | **0.0220** | 2.259 | **0.0245** | 2.294 | **0.0224** | 2.090 | **0.0374** | 2.062 | **0.0401** | 1.913 | 0.0566 | **Average FD** |
| **Scanner*** | 1.536 | 0.178 | 1.428 | 0.213 | 1.379 | 0.232 | 1.174 | 0.322 | 1.179 | 0.319 | 1.251 | 0.285 | 1.277 | 0.274 | **Scanner*** |
| **Mentalizing** | | | -4.417 | **1.354e-05** | -4.473 | **1.058e-05** | -1.774 | **0.0770** | -1.091 | 0.276 | -0.918 | 0.359 | -0.486 | 0.522 | **Mentalizing** |
| **IRI Empathic Concern** | | | | | 0.743 | 0.458 | 0.850 | 0.396 | 0.929 | 0.354 | 0.824 | 0.411 | 0.867 | 0.386 | **IRI Empathic Concern** |
| **MCCB Composite Score** | | | | | | | -3.069 | **0.00232** | -2.571 | **0.0106** | -2.321 | **0.0209** | -2.340 | **0.0199** | **MCCB Composite Score** |
| **BSFS 1** | | | | | | | | | 2.087 | **0.0377** | 2.124 | **0.0344** | 2.213 | **0.0276** | **BSFS 1** |
| **BSFS 2** | | | | | | | | | -1.171 | 0.242 | -1.032 | 0.303 | -0.964 | 0.336 | **BSFS 2** |
| **BSFS 3** | | | | | | | | | -1.769 | 0.0778 | -1.431 | 0.153 | -1.615 | 0.107 | **BSFS 3** |
| **BSFS 4** | | | | | | | | | 0.319 | 0.750 | 0.279 | 0.780 | 0.717 | 0.474 | **BSFS 4** |
| **BSFS 5** | | | | | | | | | 0.208 | 0.835 | 0.249 | 0.803 | 0.158 | 0.875 | **BSFS 5** |
| **BSFS 6** | | | | | | | | | -1.026 | 0.306 | -0.872 | 0.384 | -0.899 | 0.369 | **BSFS 6** |
| **Diagnosis** | | | | | | | | | | | -0.807 | 0.420 | 0.846 | 0.425 | **Diagnosis** |
| **Age*Sex** | | | | | | | | | | | | | 0.480 | 0.631 | **Age*Sex** |
| **Age*Diagnosis** | | | | | | | | | | | | | -1.854 | 0.0647 | **Age*Diagnosis** |
| **Diagnosis*Mentalizing** | | | | | | | | | | | | | 0.488 | 0.626 | **Diagnosis*Mentalizing** |
| **Diagnosis*Sex** | | | | | | | | | | | | | 0.880 | 0.379 | **Diagnosis*Sex** |
| **Mentalizing*Sex** | | | | | | | | | | | | | -0.024 | 0.981 | **Mentalizing*Sex** |
| **Adjusted $R^2$** | 0.08367 | | 0.1314 | | 0.1302 | | 0.1515 | | 0.1611 | | 0.1602 | | 0.1617 | | |
| **Adjusted $\Delta R^2$** | - | | 0.04773 | | -0.0012 | | 0.0213 | | 0.0096 | | -0.0009 | | 0.0015 | | |
| **Model Significance**** | - | | **1.354e-05** | | 0.4578 | | **0.002322** | | 0.1366 | | 0.4205 | | 0.3535 | | |

FD= framewise displacement, MCCB= MATRICS Consensus Cognitive Battery, BSFS= Birchwood Social Functioning Score, BSFS 1= Social Engagement/Withdrawal, BSFS 2= Interpersonal, BSFS 3= Independence - compliance, BSFS 4= Independence – performance, BSFS 5= Recreation, BSFS 6= Pro-social, IRI = Social Cognitive Interpersonal Reactivity Index, **Scanner*= coded as a factor, reporting the F-value alternatively, ***Model significance* = ANOVA of the delta R-squared comparing the two models.

#### Table S5: Significant Predictor Variables in Final Model of RS Hierarchical Regression

| Predictor Variable | β coefficient | F-value | P-value |
|---|---|---|---|
| Age | 0.0002516 | 3.8071 | 0.0519 |
| Sex | -0.004617 | 0.6466 | 0.422 |
| Average FD | 0.01776 | 3.6611 | 0.0566 |
| Scanner | 0.001959 | 1.2769 | 0.274 |
| Diagnostic group | 0.004262 | 0.6377 | 0.425 |
| Mentalizing | -0.0009094 | 0.4106 | 0.522 |
| IRI Empathic Concern | 0.0001338 | 0.7522 | 0.386 |
| **MCCB Composite Score** | -0.0001512 | 5.4745 | **0.0199** |
| **BSFS Social Engagement/Withdrawal** | 0.0007727 | 4.8989 | **0.0276** |
| BSFS Interpersonal | -0.0001650 | 0.9292 | 0.336 |
| BSFS Independence-compliance | -0.0001280 | 2.6079 | 0.107 |
| BSFS Independence-performance | 0.00008585 | 0.5147 | 0.474 |
| BSFS Recreation | 0.00002640 | 0.0249 | 0.875 |
| BSFS Pro-social | -0.0002517 | 0.8079 | 0.369 |
| Age*sex | 0.00006631 | 0.2305 | 0.631 |
| Age*diagnostic group | -0.0002527 | 3.4369 | 0.0647 |
| Diagnostic group*mentalizing | 0.0009540 | 0.2380 | 0.626 |
| Diagnostic group*sex | 0.002839 | 0.7747 | 0.379 |
| Mentalizing*sex | -0.00004787 | 0.0006 | 0.981 |

FD= framewise displacement, MCCB= MATRICS Consensus Cognitive Battery, BSFS= Birchwood Social Functioning Score, IRI = Social Cognitive Interpersonal Reactivity Index

### Table S6: EA Task Hierarchical Regression Including Medication Effects and Clinical Scores (SSD Only)

| | Model 1 | | Model 2 | | Model 3 | | Model 4 | | Model 5 | | Model 6 | | Model 7 | | Model 8 | | Model 9 | | |
|---|---|---|---|---|---|---|---|---|---|---|---|---|---|---|---|---|---|---|---|
| | | | Higher-level Social Cognition | | Emotional Empathy | | Neurocognition | | Functional Outcome | | Medication Effects | | Negative Symptoms | | Positive Symptoms | | Interaction Terms | | |
| | t | p | t | p | t | p | t | p | t | p | t | p | t | p | t | p | t | p | |
| **Age** | 3.176 | **0.00178** | 2.724 | **0.00715** | 2.647 | **0.00891** | 2.656 | **0.00870** | 2.559 | **0.0114** | 2.601 | **0.0102** | 2.603 | **0.0102** | 2.616 | **0.00982** | 0.743 | **0.0104** | **Age** |
| **Sex** | -0.707 | 0.481 | -0.714 | 0.477 | -0.284 | 0.777 | -0.397 | 0.692 | -0.547 | 0.585 | -0.394 | 0.694 | -0.396 | 0.692 | -0.476 | 0.634 | -0.684 | 0.641 | **Sex** |
| **Average FD** | 2.206 | **0.0288** | 2.001 | **0.0471** | 1.898 | 0.0594 | 1.984 | 0.0489 | 2.116 | **0.0360** | 2.138 | 0.034 | 2.223 | **0.0277** | 2.205 | **0.0290** | 2.429 | **0.0164** | **Average FD** |
| **Scanner*** | 0.473 | 0.796 | 0.5436 | 0.743 | 0.640 | 0.669 | 0.725 | 0.606 | 0.929 | 0.464 | 0.990 | 0.426 | 0.800 | 0.552 | 0.790 | 0.559 | 0.577 | 0.718 | **Scanner*** |
| **Mentalizing** | | | -1.608 | 0.110 | -1.955 | 0.0523 | -0.741 | 0.460 | -0.272 | 0.786 | -0.383 | 0.702 | 0.068 | 0.946 | 0.172 | 0.864 | 0.216 | 0.872 | **Mentalizing** |
| **IRI Empathic Concern** | | | | | 2.199 | **0.0292** | 2.338 | **0.0206** | 2.112 | **0.0363** | 2.170 | **0.0315** | 2.136 | **0.0343** | 2.099 | **0.037** | 1.794 | 0.0749 | **IRI Empathic Concern** |
| **MCCB Composite Score** | | | | | | | -1.740 | 0.0838 | -1.952 | 0.0528 | -1.994 | **0.0479** | -1.934 | 0.054 | -1.960 | 0.0518 | -1.997 | **0.0477** | **MCCB Composite Score** |
| **BSFS 1** | | | | | | | | | 1.165 | 0.246 | 1.173 | 0.242 | 1.434 | 0.154 | 1.290 | 0.199 | 1.090 | 0.277 | **BSFS 1** |
| **BSFS 2** | | | | | | | | | -0.875 | 0.383 | -0.902 | 0.369 | -1.099 | 0.274 | -1.168 | 0.245 | -0.868 | 0.387 | **BSFS 2** |
| **BSFS 3** | | | | | | | | | 1.339 | 0.182 | 1.288 | 0.200 | 1.248 | 0.214 | 1.210 | 0.228 | 1.167 | 0.245 | **BSFS 3** |
| **BSFS 4** | | | | | | | | | -0.310 | 0.757 | -0.358 | 0.721 | -0.610 | 0.543 | -0.570 | 0.570 | -0.529 | 0.598 | **BSFS 4** |
| **BSFS 5** | | | | | | | | | 0.313 | 0.754 | 0.277 | 0.782 | 0.398 | 0.692 | 0.490 | 0.625 | 0.264 | 0.792 | **BSFS 5** |
| **BSFS 6** | | | | | | | | | -1.222 | 0.223 | -1.187 | 0.237 | -1.099 | 0.273 | -1.152 | 0.251 | -1.146 | 0.254 | **BSFS 6** |
| **CPZ equivalents** | | | | | | | | | | | -0.673 | 0.502 | -0.788 | 0.432 | -0.648 | 0.518 | -0.743 | 0.459 | **CPZ equivalents** |
| **SANS Affective Flattening** | | | | | | | | | | | | | 0.474 | 0.636 | 0.432 | 0.666 | 0.388 | 0.699 | **SANS Affective Flattening** |
| **SANS Alogia** | | | | | | | | | | | | | 0.975 | 0.331 | 0.921 | 0.358 | 0.944 | 0.347 | **SANS Alogia** |
| **SANS Avolition Apathy** | | | | | | | | | | | | | 1.002 | 0.318 | 1.081 | 0.281 | 0.774 | 0.440 | **SANS Avolition Apathy** |
| **SANS Anhedonia Asociality** | | | | | | | | | | | | | -1.248 | 0.214 | -1.219 | 0.225 | -1.177 | 0.241 | **SANS Anhedonia Asociality** |
| **BPRS Positive Symptoms** | | | | | | | | | | | | | | | -0.275 | 0.784 | 0.007 | 0.994 | **BPRS Positive Symptoms** |
| **BPRS Anxiety/Depression** | | | | | | | | | | | | | | | -0.486 | 0.628 | -0.354 | 0.724 | **BPRS Anxiety/Depression** |
| **Age*Sex** | | | | | | | | | | | | | | | | | 0.731 | 0.466 | **Age*Sex** |
| **Age*Mentalizing** | | | | | | | | | | | | | | | | | 1.211 | 0.228 | **Age*Mentalizing** |
| **Sex*Mentalizing** | | | | | | | | | | | | | | | | | -0.930 | 0.354 | **Sex*Mentalizing** |
| **Adjusted $R^2$** | 0.1313 | | 0.1396 | | 0.1592 | | 0.1696 | | 0.1675 | | 0.1646 | | 0.1608 | | 0.1520 | | 0.1493 | | |
| **Adjusted $\Delta R^2$** | - | | 0.0083 | | 0.0196 | | 0.0104 | | -0.0021 | | -0.0029 | | -0.0038 | | -0.0088 | | -0.0027 | | |
| **Model Significance**** | - | | 0.1098 | | **0.0293** | | 0.08384 | | 0.4728 | | 0.502 | | 0.5114 | | 0.8072 | | 0.4705 | | |

FD= framewise displacement, MCCB= MATRICS Consensus Cognitive Battery, BSFS= Birchwood Social Functioning Score, BSFS 1= Social Engagement/Withdrawal, BSFS 2= Interpersonal, BSFS 3= Independence - compliance, BSFS 4= Independence – performance, BSFS 5= Recreation, BSFS 6= Pro-social, IRI = Social Cognitive Interpersonal Reactivity Index, SANS = Scale for the Assessment of Negative Symptoms, BPRS= Brief Psychiatric Rating Scale, **Scanner*= coded as a factor, reporting the F-value alternatively, ***Model significance* = ANOVA of the delta R-squared comparing the two models.

### Table S7: RS Hierarchical Regression Including Medication Effects and Clinical Scores (SSD Only)

| | Model 1 | | Model 2 Higher-level Social Cognition | | Model 3 Emotional Empathy | | Model 4 Neurocognition | | Model 5 Functional Outcome | | Model 6 Medication Effects | | Model 7 Negative Symptoms | | Model 8 Positive Symptoms | | Model 9 Interaction Terms | | |
|---|---|---|---|---|---|---|---|---|---|---|---|---|---|---|---|---|---|---|---|
| | t | p | t | p | t | p | t | p | t | p | t | p | t | p | t | p | t | p | |
| **Age** | 3.208 | **0.00161** | 2.590 | **0.0105** | 2.571 | **0.0110** | 2.615 | **0.0098** | 2.296 | **0.0230** | 2.258 | **0.0254** | 1.547 | 0.124 | 1.537 | 0.127 | 0.451 | 0.133 | **Age** |
| **Sex** | -1.501 | 0.135 | -1.519 | 0.131 | -1.478 | 0.141 | -1.685 | 0.0940 | -1.477 | 0.142 | -1.499 | 0.136 | -1.342 | 0.182 | -1.331 | 0.185 | -0.729 | 0.189 | **Sex** |
| **Average FD** | 0.547 | 0.585 | 0.613 | 0.541 | 0.610 | 0.543 | 0.850 | 0.397 | 0.726 | 0.469 | 0.688 | 0.493 | 0.784 | 0.434 | 0.784 | 0.435 | 0.842 | 0.401 | **Average FD** |
| **Scanner*** | 1.642 | 0.152 | 1.444 | 0.211 | 1.401 | 0.227 | 1.372 | 0.238 | 1.106 | 0.360 | 1.096 | 0.365 | 1.018 | 0.409 | 1.003 | 0.418 | 0.948 | 0.452 | **Scanner*** |
| **Mentalizing** | | | -2.114 | **0.0360** | -2.094 | **0.0378** | -0.201 | 0.841 | 0.500 | 0.617 | 0.542 | 0.589 | 0.592 | 0.555 | 0.592 | 0.555 | 0.400 | 0.565 | **Mentalizing** |
| **IRI Empathic Concern** | | | | | 0.048 | 0.962 | 0.299 | 0.765 | 0.126 | 0.900 | 0.093 | 0.926 | 0.033 | 0.974 | 0.020 | 0.984 | -0.143 | 0.887 | **IRI Empathic Concern** |
| **MCCB Composite Score** | | | | | | | -3.053 | **0.00266** | -3.355 | **0.00100** | -3.308 | **0.00117** | -3.256 | **0.00140** | -3.164 | **0.00189** | -3.162 | **0.00191** | **MCCB Composite Score** |
| **BSFS 1** | | | | | | | | | 2.895 | **0.00435** | 2.877 | **0.00459** | 2.921 | **0.00403** | 2.848 | **0.00504** | 2.655 | **0.00883** | **BSFS 1** |
| **BSFS 2** | | | | | | | | | -0.203 | 0.839 | -0.188 | 0.851 | 0.175 | 0.861 | 0.155 | 0.877 | 0.260 | 0.795 | **BSFS 2** |
| **BSFS 3** | | | | | | | | | -0.871 | 0.385 | -0.849 | 0.397 | -0.741 | 0.460 | -0.739 | 0.461 | -0.738 | 0.462 | **BSFS 3** |
| **BSFS 4** | | | | | | | | | -0.375 | 0.709 | -0.349 | 0.728 | -0.395 | 0.693 | -0.379 | 0.705 | -0.328 | 0.743 | **BSFS 4** |
| **BSFS 5** | | | | | | | | | 0.486 | 0.628 | 0.505 | 0.614 | 0.937 | 0.350 | 0.932 | 0.353 | 0.752 | 0.453 | **BSFS 5** |
| **BSFS 6** | | | | | | | | | -1.899 | 0.0595 | -1.907 | 0.0583 | -1.820 | 0.0708 | -1.807 | 0.0728 | -1.766 | 0.0796 | **BSFS 6** |
| **CPZ equivalents** | | | | | | | | | | | 0.281 | 0.779 | 0.242 | 0.809 | 0.254 | 0.800 | 0.159 | 0.874 | **CPZ equivalents** |
| **SANS Affective Flattening** | | | | | | | | | | | | | -0.953 | 0.342 | -0.946 | 0.346 | -0.964 | 0.337 | **SANS Affective Flattening** |
| **SANS Alogia** | | | | | | | | | | | | | 0.663 | 0.508 | 0.654 | 0.514 | 0.656 | 0.513 | **SANS Alogia** |
| **SANS Avolition Apathy** | | | | | | | | | | | | | 0.930 | 0.354 | 0.925 | 0.356 | 0.747 | 0.456 | **SANS Avolition Apathy** |
| **SANS Anhedonia Asociality** | | | | | | | | | | | | | 0.673 | 0.502 | 0.666 | 0.506 | 0.645 | 0.520 | **SANS Anhedonia Asociality** |
| **BPRS Positive Symptoms** | | | | | | | | | | | | | | | -0.081 | 0.936 | 0.047 | 0.963 | **BPRS Positive Symptoms** |
| **BPRS Anxiety/Depression** | | | | | | | | | | | | | | | -0.015 | 0.987 | 0.041 | 0.967 | **BPRS Anxiety/Depression** |
| **Age*Sex** | | | | | | | | | | | | | | | | | 0.451 | 0.653 | **Age*Sex** |
| **Age*Mentalizing** | | | | | | | | | | | | | | | | | 0.584 | 0.560 | **Age*Mentalizing** |
| **Sex*Mentalizing** | | | | | | | | | | | | | | | | | -0.613 | 0.541 | **Sex*Mentalizing** |
| **Adjusted $R^2$** | 0.1421 | | 0.1600 | | 0.1548 | | 0.1963 | | 0.2270 | | 0.2224 | | 0.2121 | | 0.2014 | | 0.1894 | | |
| **Adjusted $\Delta R^2$** | - | | 0.0179 | | -0.0052 | | 0.0415 | | 0.0307 | | -0.0046 | | -0.0103 | | -0.0107 | | -0.0120 | | |
| **Model Significance**** | - | | **0.03604** | | 0.9619 | | **0.002658** | | 0.06104 | | 0.7791 | | 0.7357 | | 0.9960 | | 0.8417 | | |

FD= framewise displacement, MCCB= MATRICS Consensus Cognitive Battery, BSFS= Birchwood Social Functioning Score, BSFS 1= Social Engagement/Withdrawal, BSFS 2= Interpersonal, BSFS 3= Independence - compliance, BSFS 4= Independence – performance, BSFS 5= Recreation, BSFS 6= Pro-social, IRI = Social Cognitive Interpersonal Reactivity Index, SANS = Scale for the Assessment of Negative Symptoms, BPRS= Brief Psychiatric Rating Scale, **Scanner*= coded as a factor, reporting the F-value alternatively, ***Model significance* = ANOVA of the delta R-squared comparing the two models.

### Figure 1: Q-Q plots of EA task & Resting State Hierarchical Regression Models Residuals

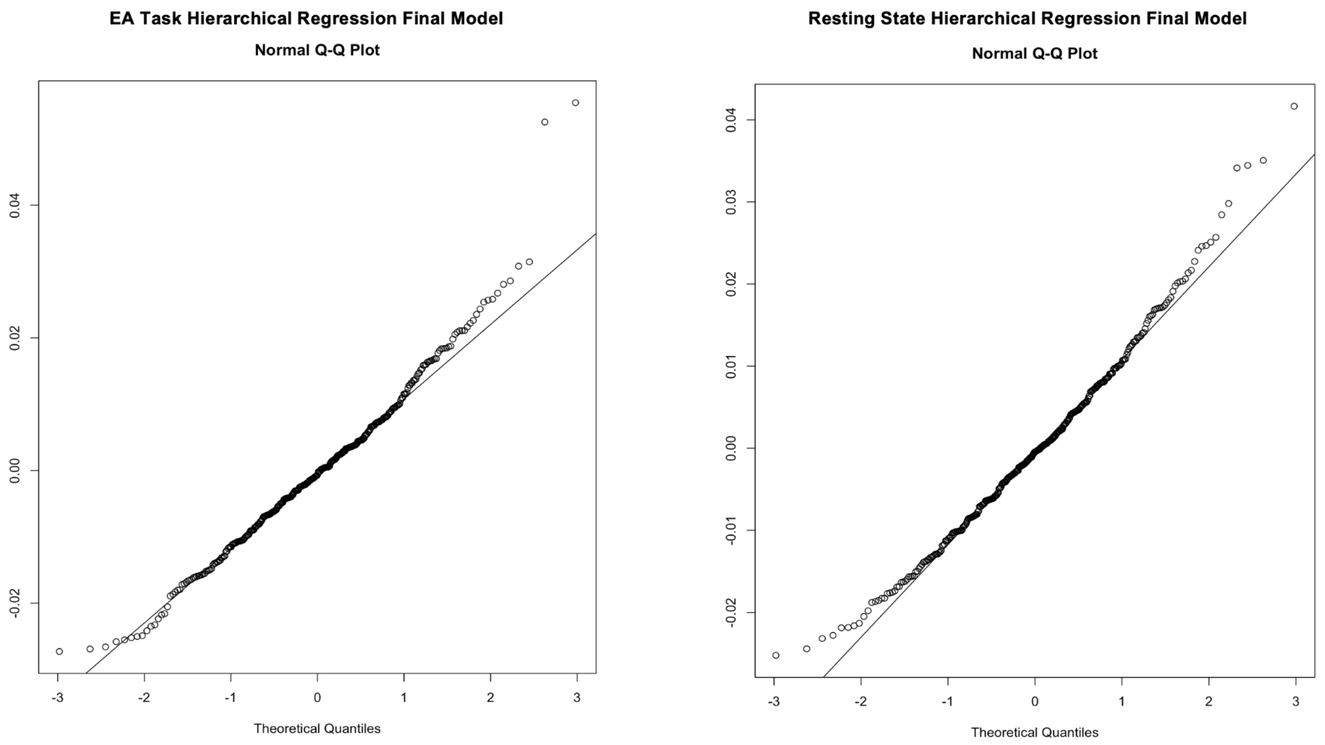
